## Supplementary Text for "The smfBox: an open-source platform for single-molecule FRET"

***Supplementary Material Contents***

| **Supplementary Information** | **Page** |
| --- | --- |
| **Supplementary Methods 1**  Hardware | **2-8** |
| **Supplementary Methods 2-**  Operational Software – C++ | **9-14** |
| **Supplementary Methods 3-**  Operational Software – LABVIEW | **15-16** |
| **Supplementary Methods 4**  Data format | **16** |
| **Supplementary Methods 5**  DNA sequences | **17** |
| **Supplementary Methods 6**  Accurate FRET validation | **18** |
| **Supplementary Results 1**  Accurate FRET results | **18** |
| **Supplementary Note 1**  Asymmetric ALEX | **19-23** |
| **Supplementary Note 2**  Verification of Dynamics | **24** |
| **Supplementary Note 3**  Possible Expansions | **25** |
| **Supplementary Equations**  1-5 | **25** |
| **Supplementary References**  22-25 | **26** |

***Supplementary Methods 1. Hardware***

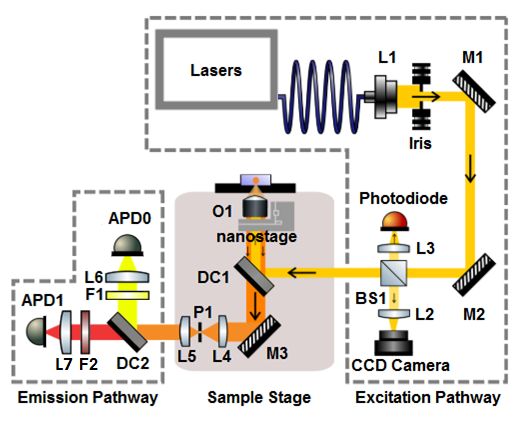
 **Supplementary Fig. 1**: 2D Schematic of the smfBox showing the Excitation pathway, box, and emission pathway

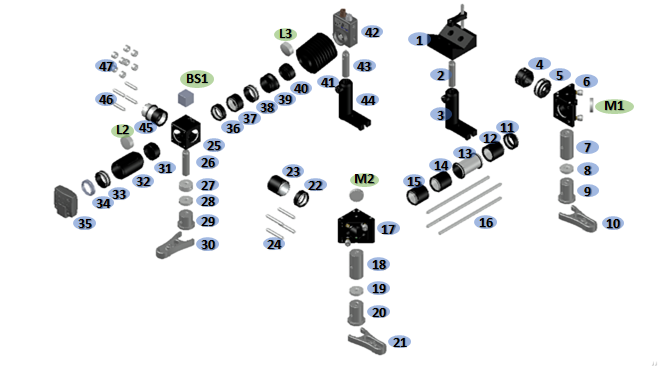
**Supplementary Fig. 2: Exploded view of the excitation path with all parts labelled according to Supplementary Table 1.**

**Supplementary Table 1:** Excitation path components

| **Part Nr** | **Product Code** | **Use** | **Part Nr** | **Product Code** | **Use** |
| --- | --- | --- | --- | --- | --- |
| 1 | VC3C/M | V-shaped collimating lens holder | 27 | RS10/M | Post spacer BS1 (L=10mm) |
| 2 | TR50/M | Collimating lens post | 28 | RS5/M | Post spacer (L=5mm) |
| 3 | UPH50/M | Post holder | 29 | RS1P8E | Pedestal Pillar Post BS1 (L=1'') |
| 4 | SM1P1 | Attachment lens to iris | 30 | CF125C/M | Clamping fork for BS1 post |
| 5 | SM1D12C | Iris | 31 | SM1L03 | Tube 1 BS1 to DCC |
| 6 | KCB1C/M | Right-Angle Kinematic Mirror Mount for M1 | 32 | SM1L20C | Slotted lens tube for L2 |
| 7 | RS50/M | Pillar post (L=50 mm) | 33 | SM1T2 | Tube coupler BS1 to DCC |
| 8 | RS5/M | Post spacer (L=5mm) | 34 | SM1A9 | Adapter SM1/C-mount BS1 to DCC |
| 9 | RS1.5P/M | Pedestal pillar post (L=38mm) | 35 | DCC1545M | CCD Camera |
| 10 | CF125C/M | Clamping fork for M1 post | 36 | SM1L03 | Tube 1 BS1 to photodetector |
| 11 | SM1L03 | Tube 1 M1 to M2 | 37 | SM1L05 | Tube 2 BS1 to photodetector |
| 12 | SM1L10 | Tube 2 M1 to M2 | 38 | SM1T2 | Tube coupler BS1 to photodetector |
| 13 | SM1T20 | Tube coupler M1 to M2 | 39 | SM1L03T | Angled tube BS1 to photodetector |
| 14 | SM1L10 | Tube 3 M1 to M2 | 40 | SM1L03 | Tube with L3 |
| 15 | SM1L10 | Tube 4 M1 to M2 | 41 | SM1B3 | Tube Bellow BS1 to photodetector |
| 16 | ER6-P4 | 4x rods M1 to M2 (6'') | 42 | DET10A/M | Photodetector |
| 17 | KCB1C/M | Right-Angle Kinematic Mirror Mount for M2 | 43 | UPH50/M | Post holder |
| 18 | RS50/M | Pillar post (L=50 mm) | 44 | TR50/M | Post for photodetector |
| 19 | RS5/M | Post spacer (L=5mm) | 45 | SM1V10 | Tube BS1 to box |
| 20 | RS1.5P/M | Pedestal pillar post (L=38mm) | 46 | ER1-P4 | 4x Cage Rods BS1 to box |
| 21 | CF125C/M | Clamping fork for M1 post | 47 | ERSCA-P4 | 2x4 Rod adapter BS1 to box |
| 22 | SM1L03 | Tube 1 M2 to BS1 | M1 | BB1-E02 | First Mirror |
| 23 | SM1S10 | Tube 2 M2 to BS1 | M2 | BB1-E02 | Second Mirror |
| 24 | ER1.5-P4 | 1x4 rods M2 to BS1 (1.5'') | BS1 | 21012 | 10/90 Beamsplitter |
| 25 | CM1-DCH/M | Cage Cube for BS1 | L2 | AC254-030-AML | Lens for Camera |
| 26 | TR50/M | Post for BS1(L=50mm) | L3 | 49793 | Lens for Photodetector |

The excitation pathway is built from a precision iris, two kinematically mounted mirrors, a beam splitter, a photodetector, and a CCD camera. Optomechanical components are mounted on non-height adjustable posts with cage-rods between them to ensure robustness of alignment. After being trimmed by the iris (Supplementary Fig. 1), the mirrors M1 and M2 give complete axial stabilisation of the beam before entering the box. The beam splitter BS1 on the first pass (blue line) reflects 10% of light through a lens (L3) onto a nanosecond rise time photodetector for calibration of laser alternation and power, whilst permitting the other 90% through to the cube. Backscattered light from the sample (green beam) is reflected off the back of the beam splitter, through a lens (L2) onto a CCD camera which is used for focusing the objective. One potential problem can arise from reflections off the photodetector passing through the beam splitter onto the CCD (dashed red line), this is averted by mounting the photodetector at a slight tilt.

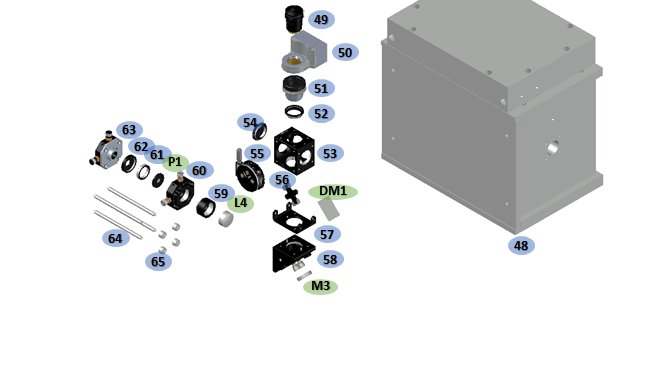

**Supplementary Fig. 3:** An exploded view of the optical components inside the microscope body, with parts labelled according to Supplementary Table 2.

**Supplementary Table 2:** Part numbered microscope body components.

| **Part Nr** | **Product Code** | **Use** | **Part Nr** | **Product Code** | **Use** |
| --- | --- | --- | --- | --- | --- |
| 48 | N/A | Machined Aluminium Box | 59 | SM1L05 | Lens Tube L4 (0.5'') |
| 49 | N1480700 | Objective | 60 | CXY1 | Translating Lens Mount for L4 |
| 50 | FOC.300 | Nanostage Z | 61 | SM1A3 | Adapter (Ext. SM1 Int. RMS) for P1 to Mount |
| 51 | N/A | Piezoconcept bushing (with nanostage) | 62 | SM1A1 | Adapter (Ext. SM05 Int. SM1) for P1 to Mount |
| 52 | SM1A12 | FOC bushing adapter (Ext. SM1 Int. M25) | 63 | CXYZ05/M | Translating Mount for P1 |
| 53 | C4W | Cage Cube DM1 | 64 | ER4-P4 | 1x4 4'' Rods |
| 54 | SM1CP2 | Cap for spare DM1 cage cube ports | 65 | ERSCA-P4 | 1x4 Rod Adapter |
| 55 | B4CRP/M | Precision mount for DM1 | DM1 | ZT532/640rpc | Excitation Dichroic |
| 56 | FFM1 | Clamp for DM1 | M3 | MM3-311-t6-1 25.4 | Hard Mirror between DM1 and L4 |
| 57 | C4W-CC | Cage Cube Connector M3 to DM1 | L4 | Edmund - 49792 | Lens before pinhole (50 mm FL) |
| 58 | KCB1/M | Right angle KM mount for M3 | P1 | PNH-20 | Pinhole (20 μm) |

Upon entering the box, lasers are reflected by the excitation dichroic DM1 into the objective and sample. The objective can be moved through z by a nanostage, and the sample can be positioned in xy if necessary (allowing for potential applications in confocal scanning techniques). Emitted light from the sample then passes back through the objective and is permitted through DM1, and onto a hard mirror M3. Lens L4 then focuses emitted light through a pinhole P1 to remove out of focus light before reaching a second lens L5 (shown in emission pathway). L4 can be adjusted in xy, P1 has full adjustment in xyz and L5 is static.

**Supplementary Fig. 4:** An exploded view of the aluminium box with components labelled according to Supplementary Table 3.

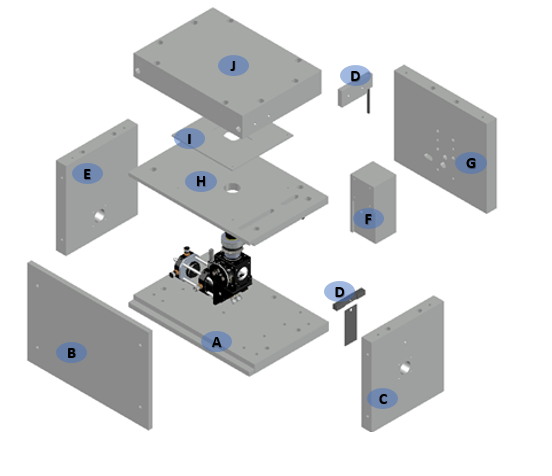

**Supplementary Table 3:** Aluminium Box components. See technical drawing in supplementary files for engineering diagrams.

| **Part Nr** | **Use** | **Part Nr** | **Use** |
| --- | --- | --- | --- |
| A | Base Plate | F | Block |
| B | Front Plate | G | Back Plate |
| C | Laser-In Plate | H | Top Plate |
| D | Shutter | I | Stage |
| E | Laser-Out Plate | J | Lid |

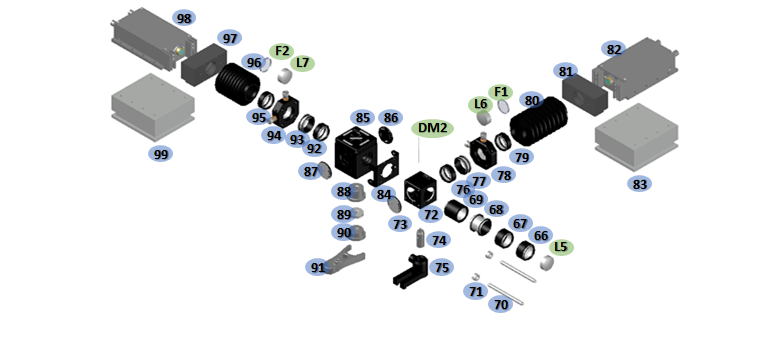

**Supplementary Fig. 5:** An exploded view of the Emission pathway with components labelled according to Supplementary Table 3.

**Supplementary Table 4:** Emission path components. Technical drawings for the APD cover and mount (81, 83, 97, 99) can be found in the supplementary files.

| **Part Nr** | **Product Code** | **Use** | **Part Nr** | **Product Code** | **Use** |
| --- | --- | --- | --- | --- | --- |
| 66 | SM1L05 | Tube 1 box to DM2 with L5 | 86 | SM1CP2 | Cap 1 for spare ports of filter cube |
| 67 | SM1L05 | Tube 2 box to DM2 | 87 | SM1CP2 | Cap 2 for spare ports of filter cube |
| 68 | SM1T10 | Tube coupler box to DM2 | 88 | RS075P/M | Pedestal pillar post filter cube (L=19mm) |
| 69 | SM1L10 | Tube 3 box to DM2 | 89 | RS5/M | Spacer (L=5mm) |
| 70 | ER3 | 2x Rods box to DM2 3'' | 90 | RS05P | Pedestal pillar post filter cube (L=0.5'') |
| 71 | ERSCA | 2x Rod adapters box to DM2 | 91 | CF125C/M | Clamping fork filter cube |
| 72 | CM1-DCH/M | Cage cube for DM2 | 92 | SM1L03 | Tube 1 DM2 to APD0 |
| 73 | SM1CP2 | Cap for spare port of DM2 cage cube | 93 | SM1T2 | Tube adapter DC2 to APD1 |
| 74 | TR30/M | Post DM2 (L=30 mm) | 94 | CXY1 | Translating lens mount L7 DC2 to APD1 |
| 75 | UPH30/M | Post Holder DM2 (L=30mm) | 95 | SM1L03 | Tube for F2 DC2 to APD1 |
| 76 | SM1L03 | Tube 1 DM2 to APD0 | 96 | SM1B3 | Lens tube bellow DC1 to APD1 |
| 77 | SM1T2 | Tube adapter DM2 to APD0 | 97 | Fabricated Al | Connector tube to APD1 |
| 78 | CXY1 | Translating lens mount L6 DM2 to APD0 | 98 | SPCM-AWRH-14 | APD1 (Excelitas) |
| 79 | SM1L03 | Tube for F1 DM2 to APD0 | 99 | Fabricated Al | Post APD1 |
| 80 | SM1B3 | Lens tube bellow DM1 to APD0 | L5 | Edmund - 49793 | Lens after Pinhole (63.5 mm FL) |
| 81 | Fabricated Al | Connector tube to APD0 | L7 | AC254-75-B-ML | APD1 Lens |
| 82 | SPCM-AWRH-14 | APD0 (Excelitas) | L6 | AC254-100-A-ML | APD0 Lens |
| 83 | Fabricated Al | Post APD0 | F2 | FF01-679/41-25 | Acceptor cleanup filter |
| 84 | CM1-CC | Cage cube connector DM2 to filter cube | F1 | FF01-571/72-25 | Donor cleanup filter |
| 85 | DFM1/M | Fluorescence filter cube | DM2 | NC395323-T640lpxr | Emission Dichroic |

The first dichroic mirror in the emission pathway, DM2, reflects light below 640 nm, sending donor emission into the path to APD0 whilst permitting acceptor photons through to APD1. In both cases a band-pass filter (F1, F2) is used to clean up laser bleed-through and Raman scatter from the emission, and a lens (L6, L7) is used to focus the beam onto the APD.

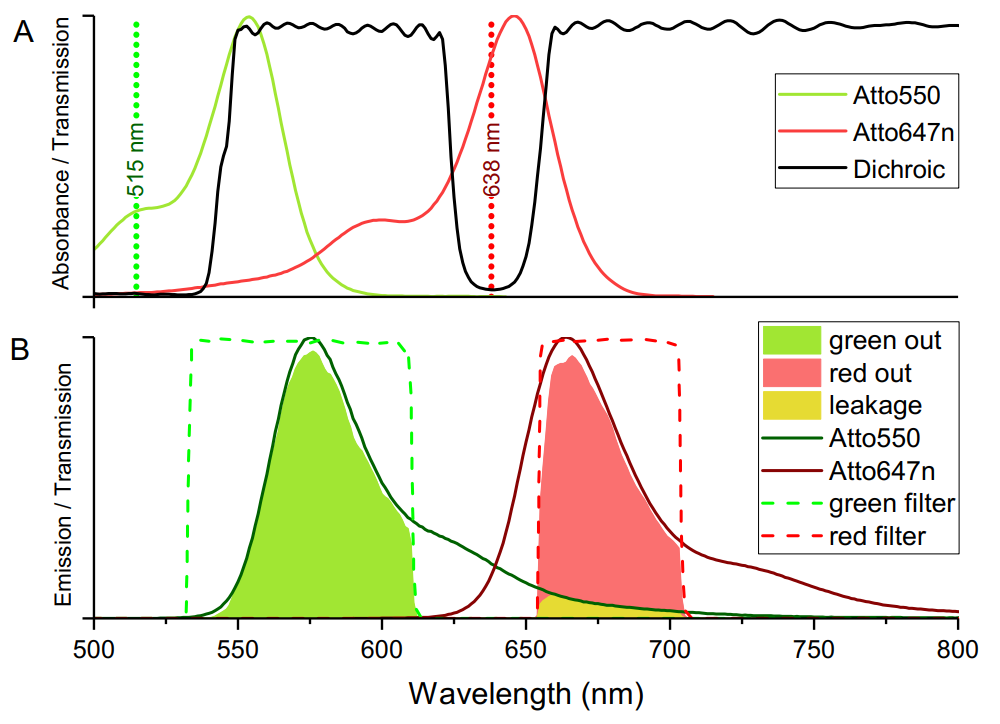
**Supplementary Fig. 6:** Spectra of optical components in the smfBox. **a:** The excitation dichroic in black with laser lines and typical dye absorption spectra in green and red. **b:** The two emission channels shown in dashed lines, with typical emission spectra. Block colours show the spectra of emission which ultimately reaches the detectors.

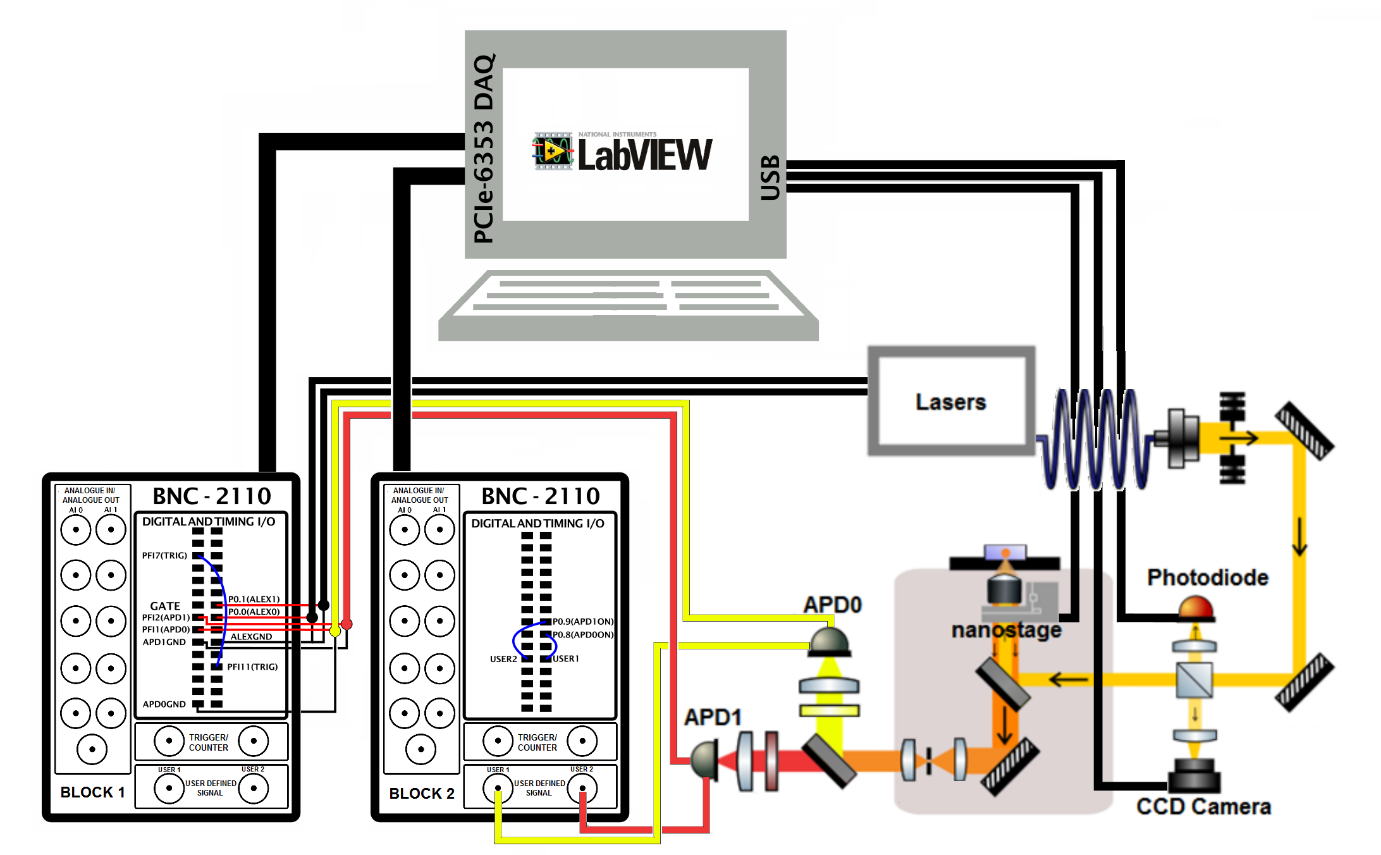

**Supplementary Fig. 7:** Wiring diagram of electronics in the smfBox. Lasers are controlled, and APD’s are gated and monitored via BNC adaptors connected to the NI-DAQ board in the PC. The CCD camera and nanostage for Z focussing, as well as the photodiode for laser power monitoring are connected directly via USB.

Once the main cube body is attached the optical table the scope can be built and aligned (~ 2 days work).

- 1. Once the main excitation pathway: M1, M2 and BS1 have been mounted they should be connected with 3 of the 4 cage rods without the lens tubes attached. The initial alignment can be done by aiming the beam to the centre of the DM1 mount port, using a target on one of the dichroic cube’s mount caps. A handy trick to give fine alignment is to rotate DM1 to be 180° to the incoming beam and place a mirror directly behind it. This mirror will reflect the bleed-through of the dichroic, creating a cavity. If the beam is flat and correctly aligned coming into the dichroic mount then the reflection of the beam should match the incoming one.
- 2. Insert the fourth rod and the lens tubes and repeat the above steps to give fine alignment.
- 3. The microscope body is then aligned without insertion of the objective, nanostage or confocal system (L4, L5 and P1): DM1 should then be aligned at 45° to reflect the beam perpendicular to the plane of the optical table (aiming to be centred on the objective’s aperture when inserted). To check this alignment long rods can be extended from the DM1 mount with the top lid of the microscope body removed. Using a two frosted glass apertures or collapsible irises, the alignment can be tuned as the beam should stay in the centre of both apertures at a far distance. Adjustment of the pitch of DM1 should correct for any deviations.
- 4. Alignment of the confocal system should then be done by inserting a mirror in the position of O1. With the laser power increased the bleed-through should be visible though the dichroic. By using long rods attached to the beam-out port of the microscope body M3 can be aligned to be flat and centred.
- 5. The confocal lenses, L4 and L5 can then be positioned by checking the beam is collimated at a long distance. This can be done by measuring the beam waist with a camera or through visual inspection and adjusting the position of the lenses relative to each other.
- 6. The pinhole can then be inserted and aligned using a power meter to track the maximum transmission.
- 7. The objective lens can then be inserted and the top plate of the box attached. Using a microscope slide with a drop of water the coarse focusing of the objective lens can be found using the methods mentioned in the operational software (see below).
- 8. Using an appropriate dye (eg. Cy3B) the pinhole alignment can then be optimised further using a power meter.
- 9. If the dye concentration is high enough the excitation pathway can then be coarsely aligned by-eye without the lens bellow attached. For fine alignment, the dye concentration or laser power can be reduced. The translation stages on the APD lenses (L6 and L7) can then be used with the APD alignment software to get exact alignment.
- 10. The lens bellows should then be attached, and the system is ready for testing.

***Supplementary Methods 2. Operational Software –* smOTTER**

We provide operational software in the form of both a standalone C++ implementation called smOTTER, and a set of LABVIEW VI’s which can be more easily customised.

**Installation**

Pre-compiled executables are available from our github or the supplementary materials of this paper. To install, simply download the zip, extract it to a location of your choosing and run the smfBoxAcquisition.exe.

The executables require the following to be installed to operate correctly:

- A copy of NI-DAQmx
- A 64-bit copy of the ThorCam software

**Compilation from Source**

A copy of Visual Studio 2017 or later and an installation of Qt 5.12 is required to compile from source. The code additionally depends on the NIDAQ C SDK which comes with an installation of DAQmx, the ThorCam C SDK which comes with an installation of the ThorCam software and a copy of the HDF5 C++ library. The HDF5 library can be compiled from source (Version 1.10.4 was used to develop the software) or a compatible compiled version can be downloaded from our GitHub.

Once all the dependencies have been installed, download the source code. The project can now be opened in Qt Creator (installed by default with Qt). Alternatively, the project can be opened in Visual Studio by installing the Qt plugin and selecting import .pro file project from the Qt VS dropdown. The code can then be compiled by selecting the build (or build and run) button in your chosen IDE. The executable should appear in %SRC_DIR%/build/Debug or %SRC_DIR%/build/Release depending on your compilation settings

**User Interface**

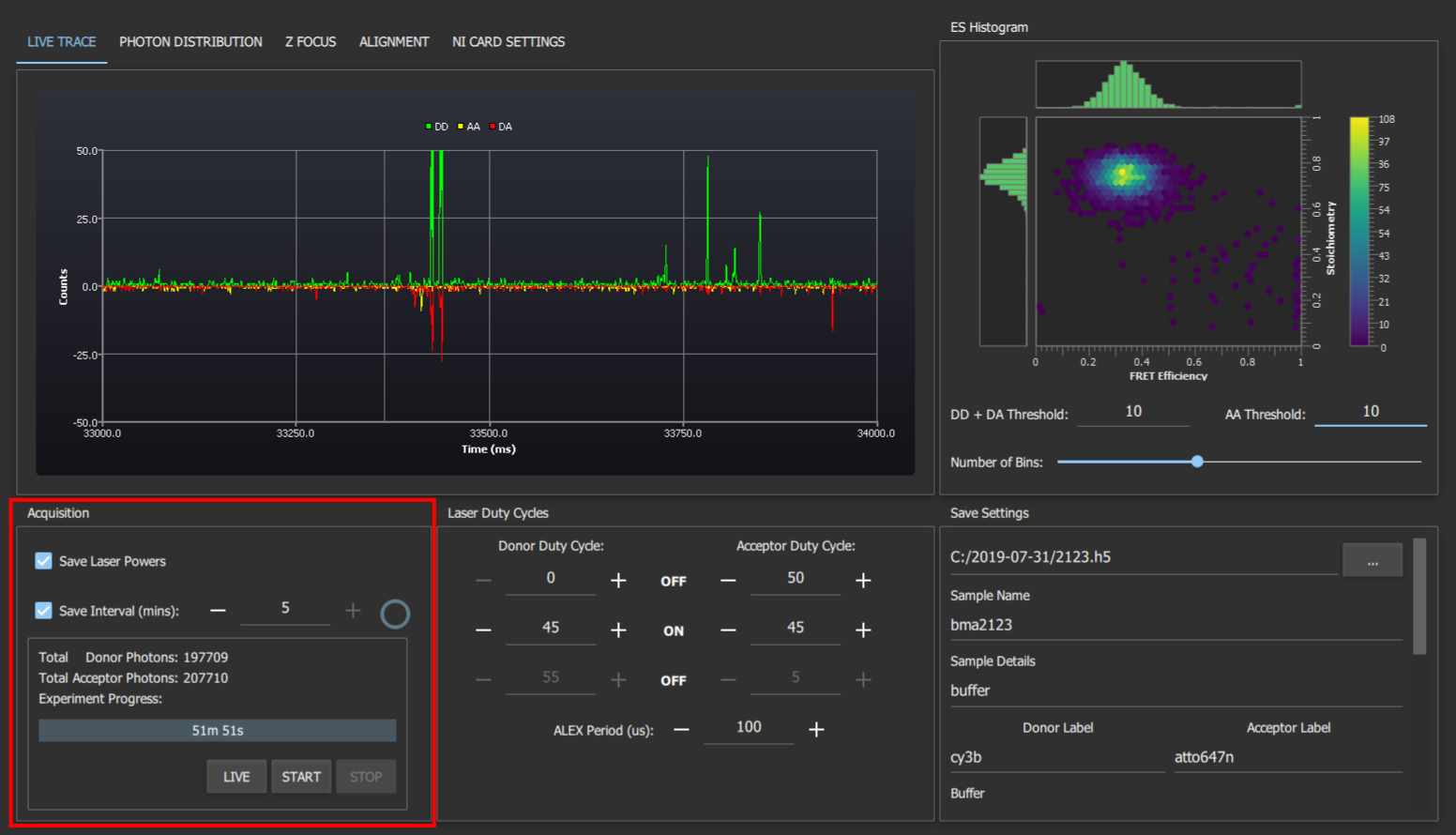

**Supplementary Fig. 8:** The acquisition settings section contains settings for and status of the current experiment. The experiment length setting controls the time the experiment should run for. The save laser powers setting can be used to perform real time recording of the laser powers as reported by the system's laser power photo-detector. The save interval checkbox toggles automatic saving data to disk during acquisition. The save interval can also be set to the desired value using the field next to the checkbox. During acquisition the circle to the right of the save interval settings indicates the time until the next save to disk. Underneath the acquisition settings is a box containing the total number of donor and acceptor photons collected, the experiment progress indicator, and the acquisition control buttons. The start and stop buttons begin and abort an acquisition respectively. The live button starts an acquisition but does not save anything to disk. This is useful for checking that everything is set up correctly before starting the acquisition.

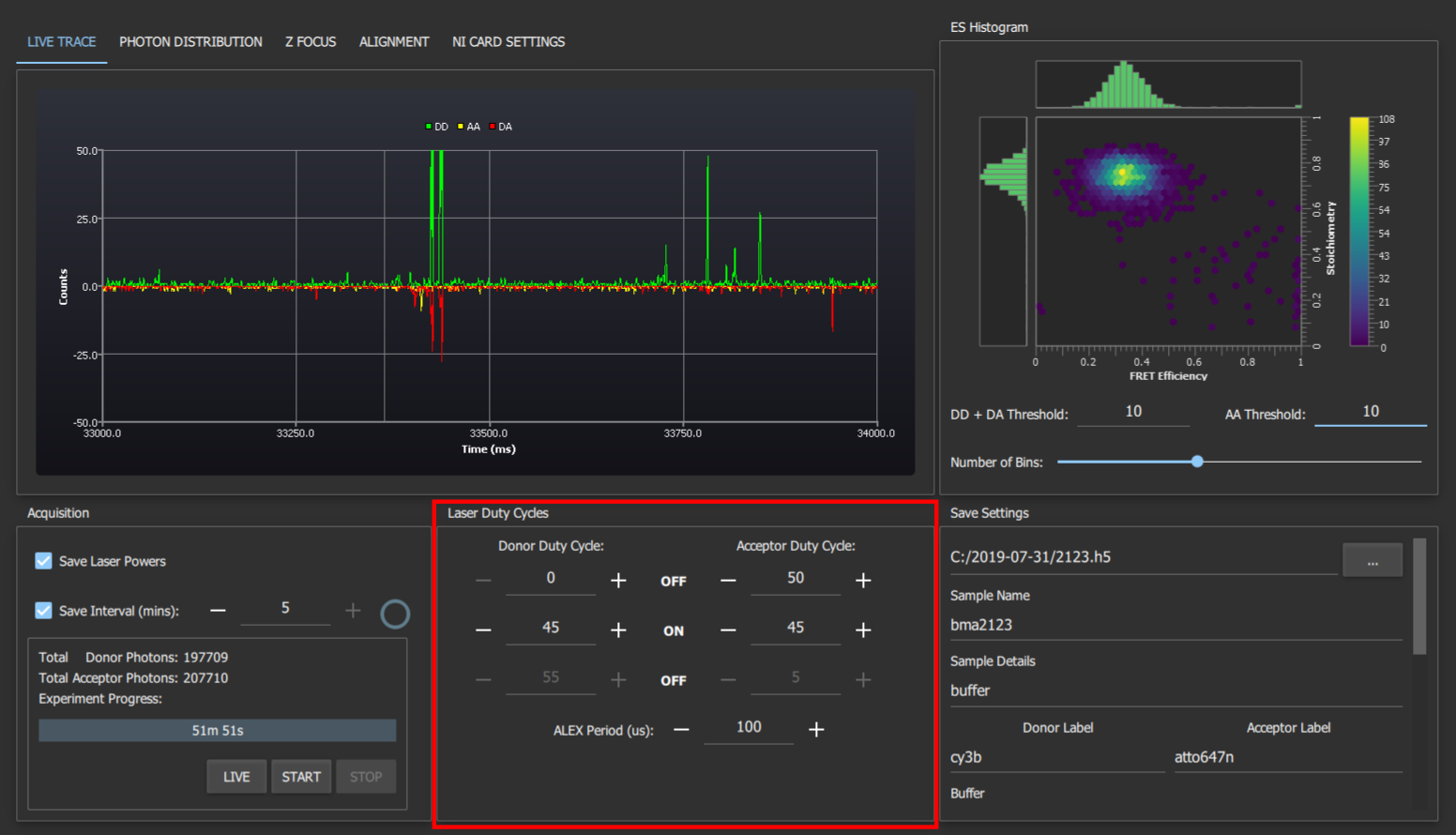

**Supplementary Fig. 9**: The laser duty cycles section allows configuration of the laser alternation periods. The first "off" row sets the time from the beginning of an ALEX period that the corresponding laser remains off for. The "on" row sets the time that the lasers remain on for. The final "off" row is the time remaining at the ALEX period that the lasers will be off for. All timings are in percentages of the ALEX period. A graphical representation of the laser duty cycles is available in the "photon distribution tab". The final setting in this section is the ALEX period in us. This is usually set to 100 us as this allows for exact synchronisation between all the functions of the NIDAQ board.

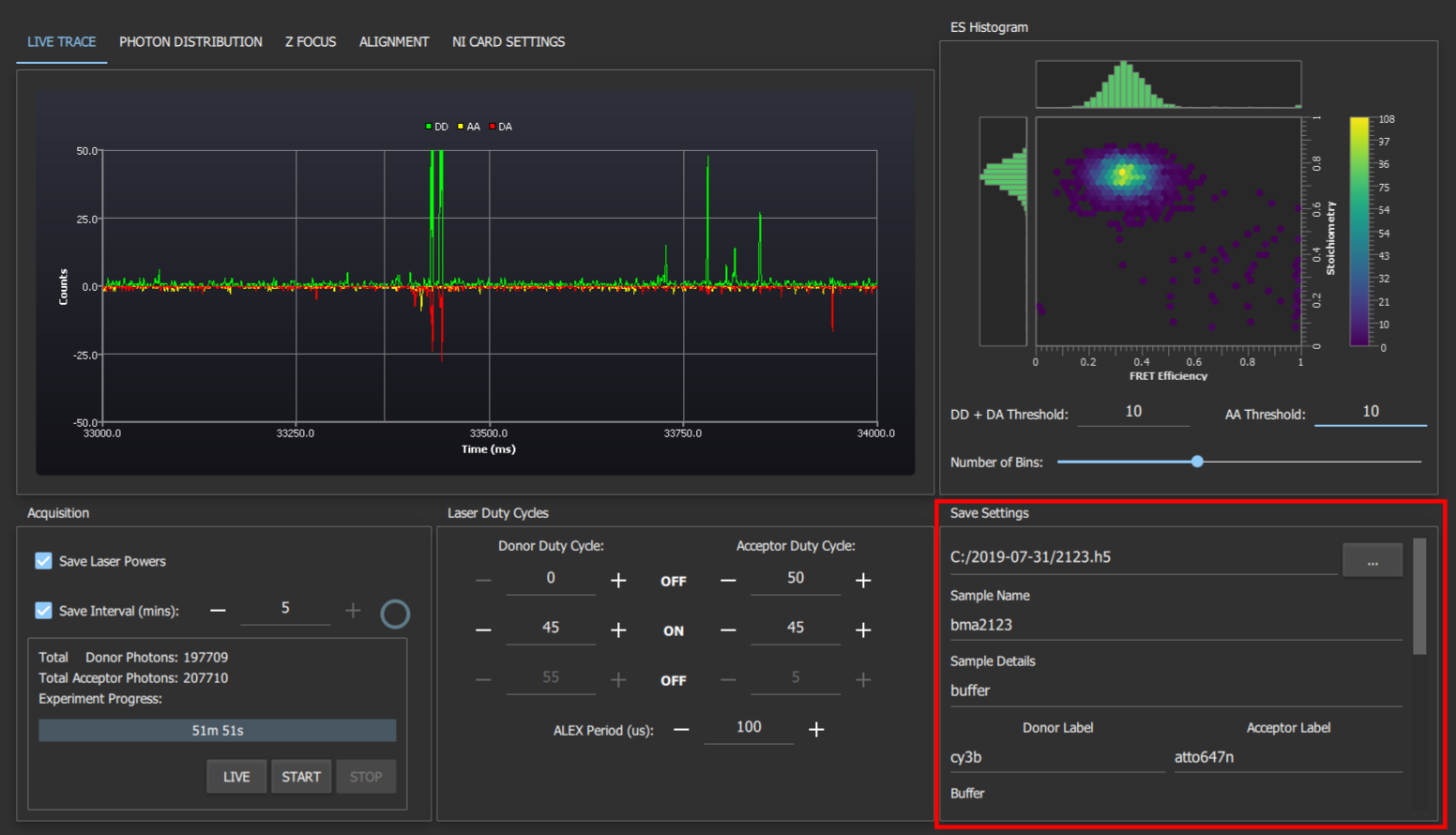

**Supplementary Fig. 10:** The save settings box contains inputs for all the metadata required by PhotonHDF5. Settings that are unlikely to change between experiments, e.g. laser wavelengths, are remembered between restarts of the program. The program will ask you before overwriting any previously saved data.

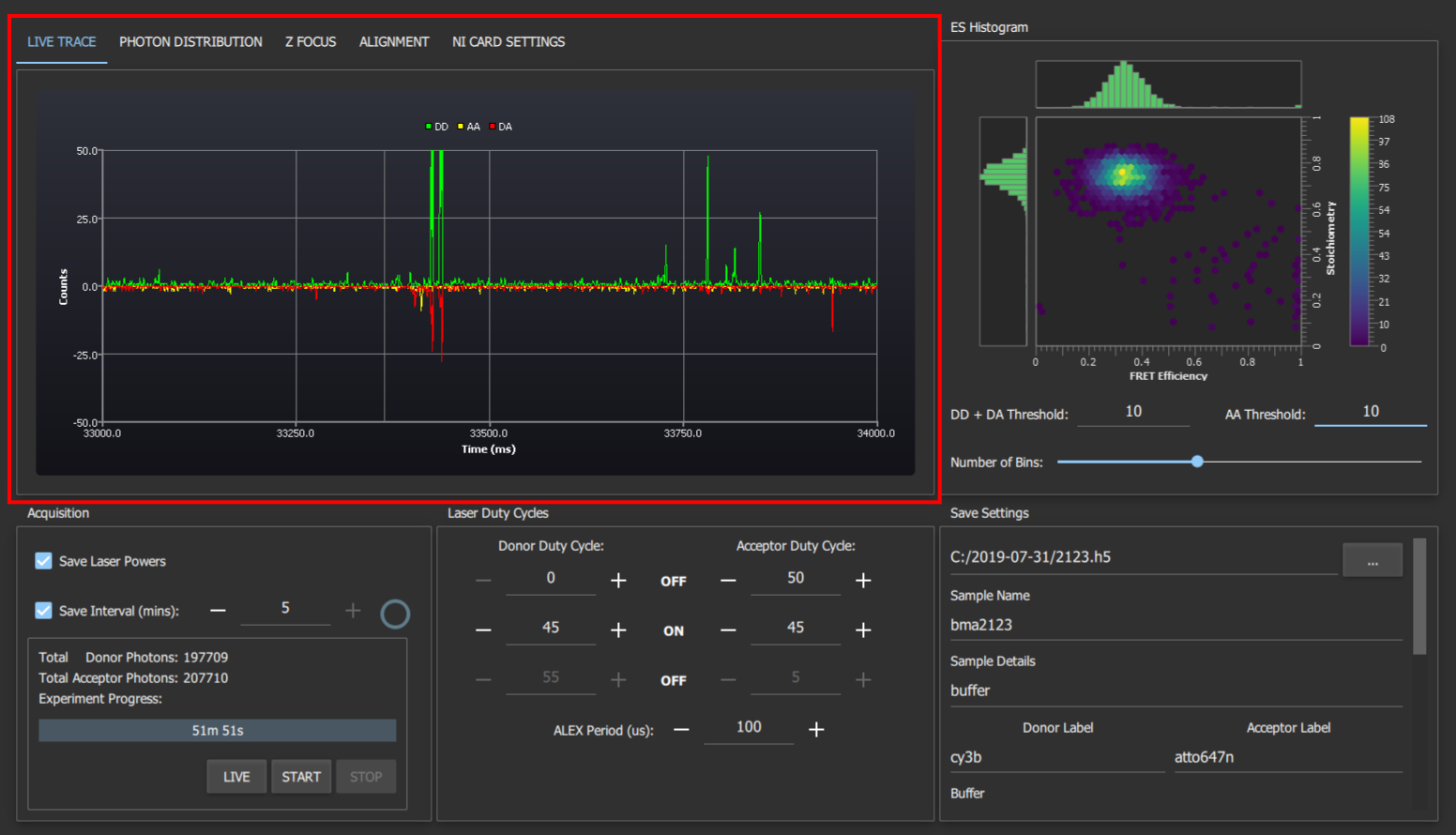

**Supplementary Fig. 11:** The live trace shows a real-time plot of the number of photons detected during live mode or an acquisition. The green trace corresponds to lasers detected on the donor APD when the donor laser is on, yellow corresponds to acceptor APD with acceptor laser and DA corresponds to acceptor APD with donor laser. The AA and DA traces are shown as inverted to make it easier to differentiate spikes in their traces from spikes in the DD trace.

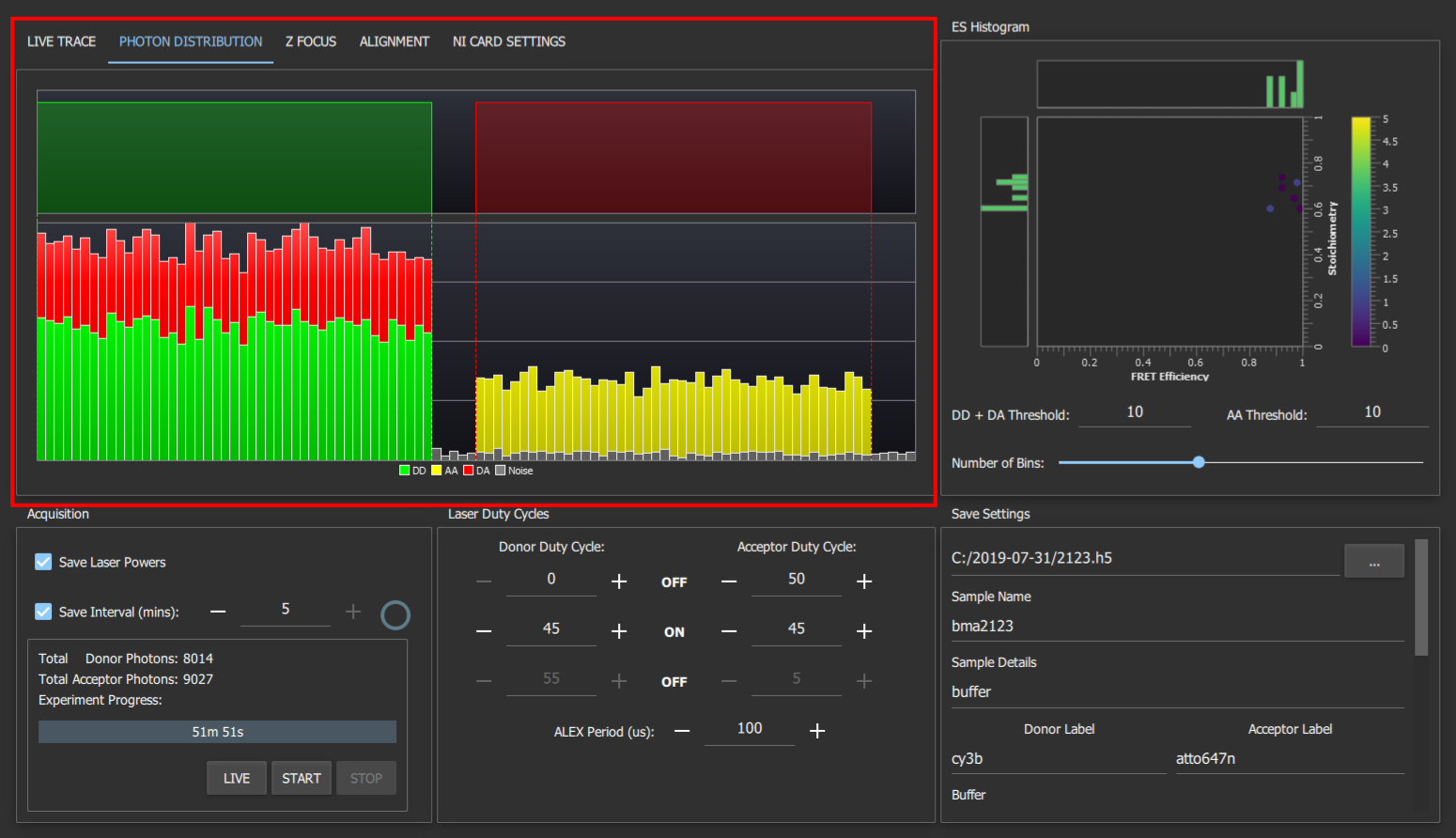

**Supplementary Fig. 12:** The photon distribution time tab shows a histogram of the photon arrival times with respect to the laser alternations (shown above the histogram). The red, green and yellow bars show the number of DA, DD and AA photons arriving at a given time relative to the laser alternation period whereas the grey bars show any detected photons that cannot be placed into one of these 3 categories (AD photons, or photons with no laser on). This panel can be a useful diagnostic to check the lasers are alternating correctly, especially if lasers with longer rise/fall times are used.

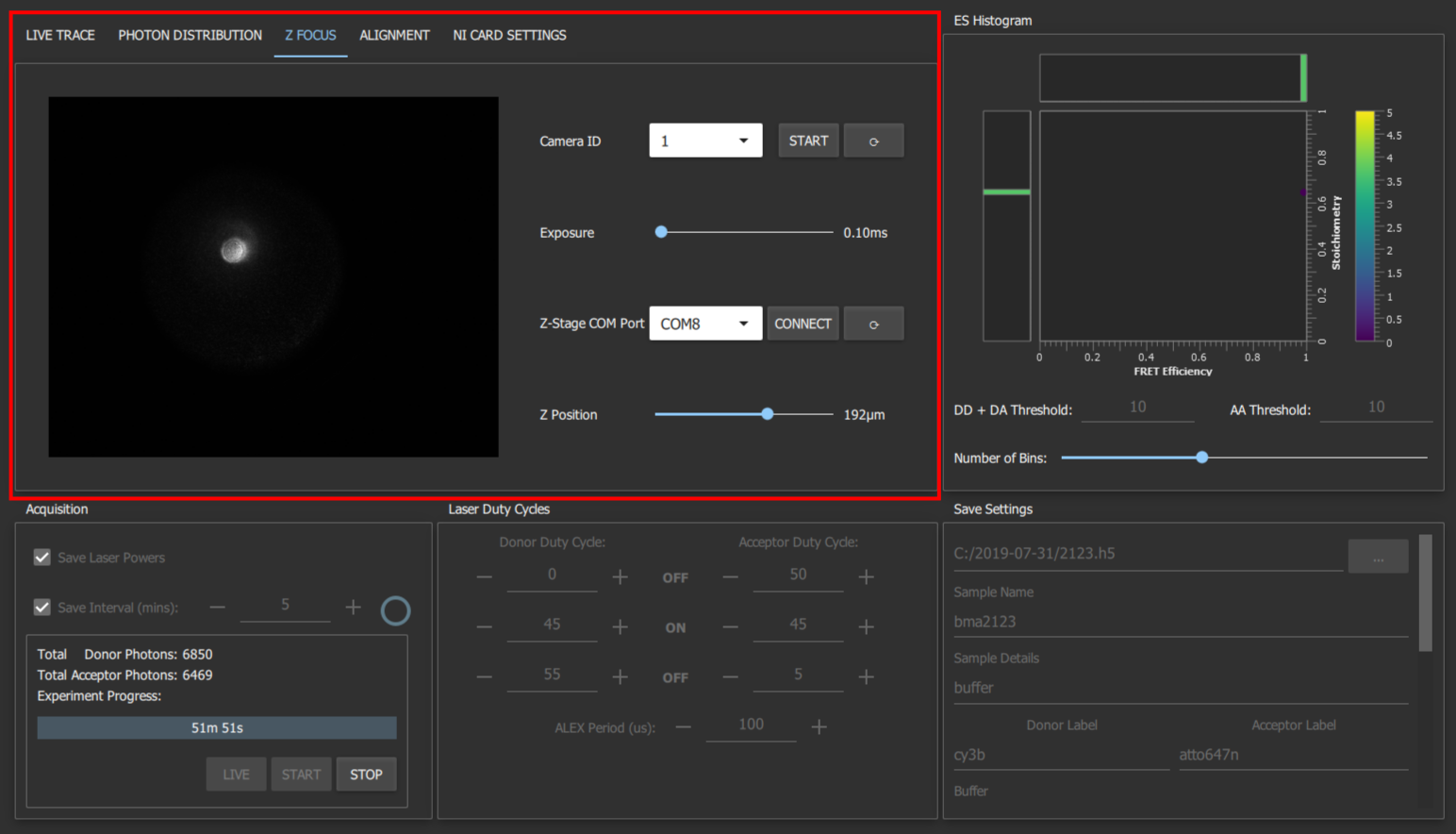

**Supplementary Fig. 13:** The Z-Focus tab allows you to control the z-position of the stage and shows the output of the CCD camera which is imaging the back-reflection of the sample. This allows you to focus the confocal spot into the sample. To operate, first place a drop of liquid on a coverslip on the objective. With either of the lasers on, move the slider for the z-position up until the back reflection shrinks to its smallest size. Then move slightly further up (~20 μm) so that the back reflection enlarges again. The confocal spot is now focused into the sample.

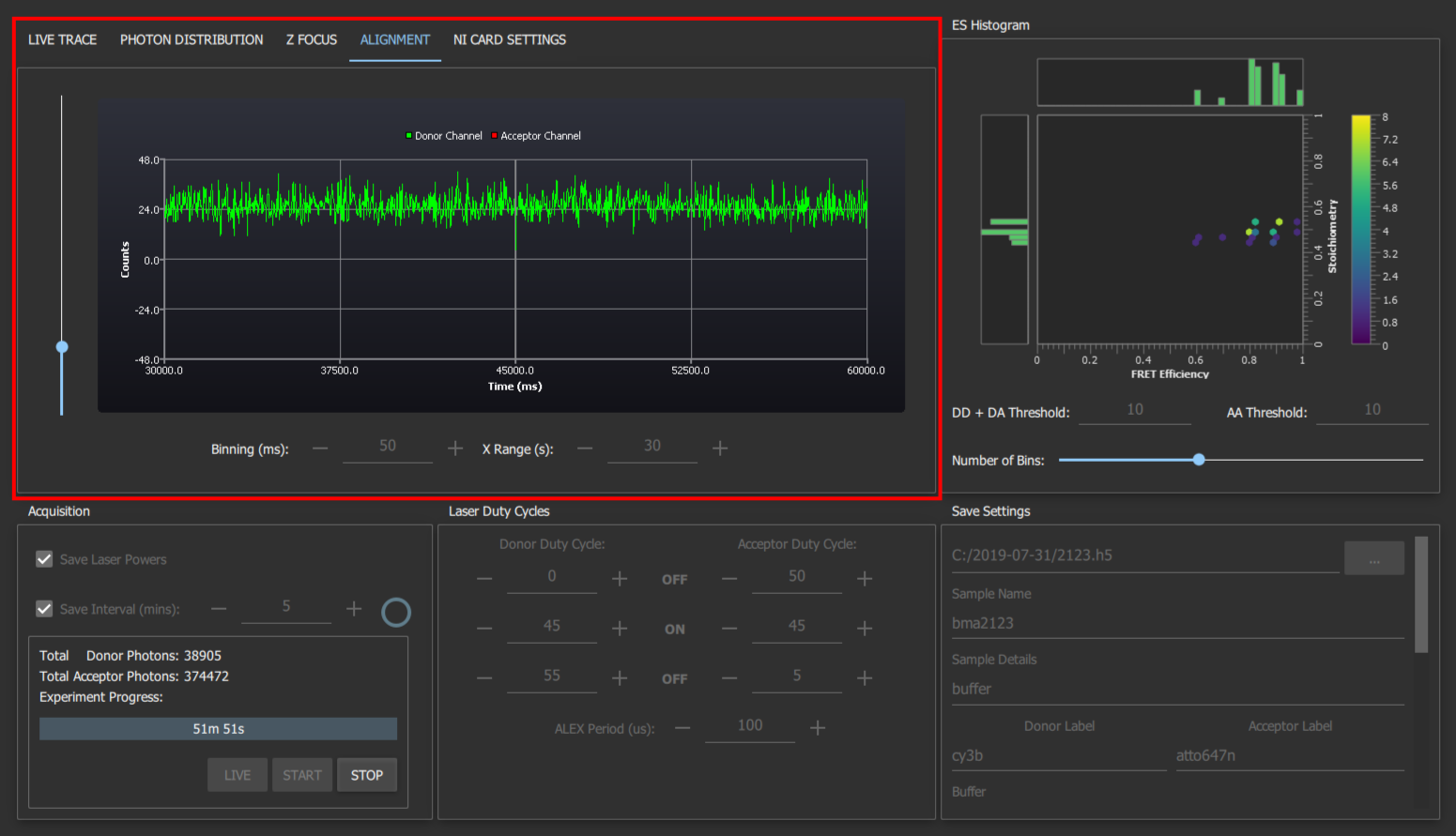

**Supplementary Fig. 14:** The alignment tab shows the number of photons detected on each detector, irrespective of which laser was on, and is able to plot a much higher flux of photons, which is useful for aligning components in the emission pathway. To make use of this, place a fluorescent sample on the coverslip which will emit in both channels when excited with the donor excitation laser (we use free Cy3B), then maximise the emission intensity reported by iteratively adjusting the positions of the APD lenses (L6 and L7), the pinhole, and if needed M3 and L4. Unlike the live trace graph, the binning and x-range can be adjusted. This allows you to select settings that result in a sufficiently smooth trace to allow you to easily align the system. For fine alignment it is recommended to use a high concentration so as to avoid fluctuations due to molecular diffusion, but low laser power such that the software can handle the signal from the APDs.

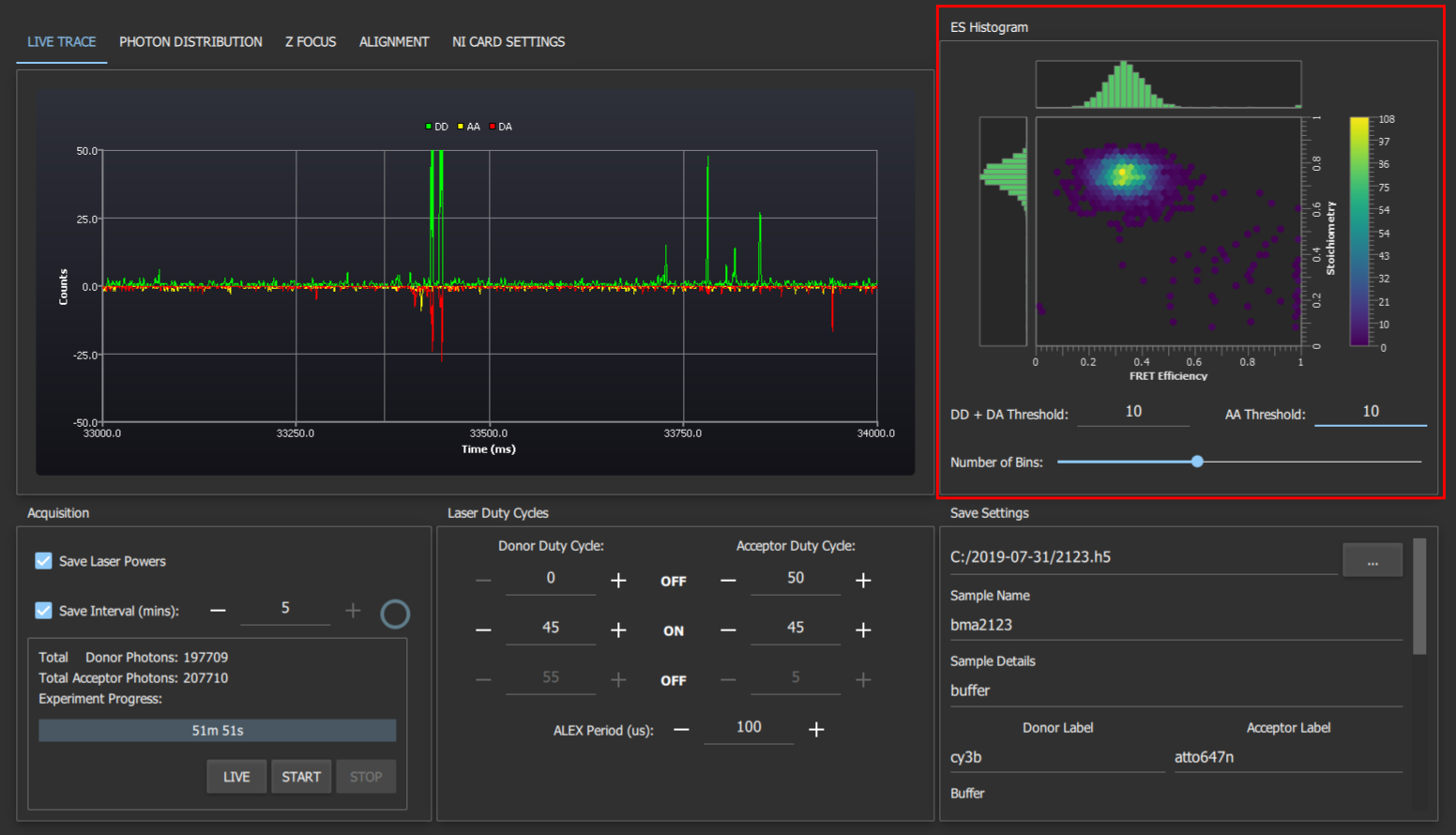

**Supplementary Fig. 15:** The live ES histogram shows an approximation of the hexagonal ES plots traditionally produced during ALEX-FRET analysis and helps give you a feel for any issues with your sample before you analyse the data. The FRET efficiency and stoichiometry are calculated and plotted for bursts which surpass both thresholds in a 1ms window. The number of bins slider can be adjusted to change the resolution of the histogram. Increasing it to the maximum value produces a plot that's more like a scatter graph if you prefer this to the hex plot.

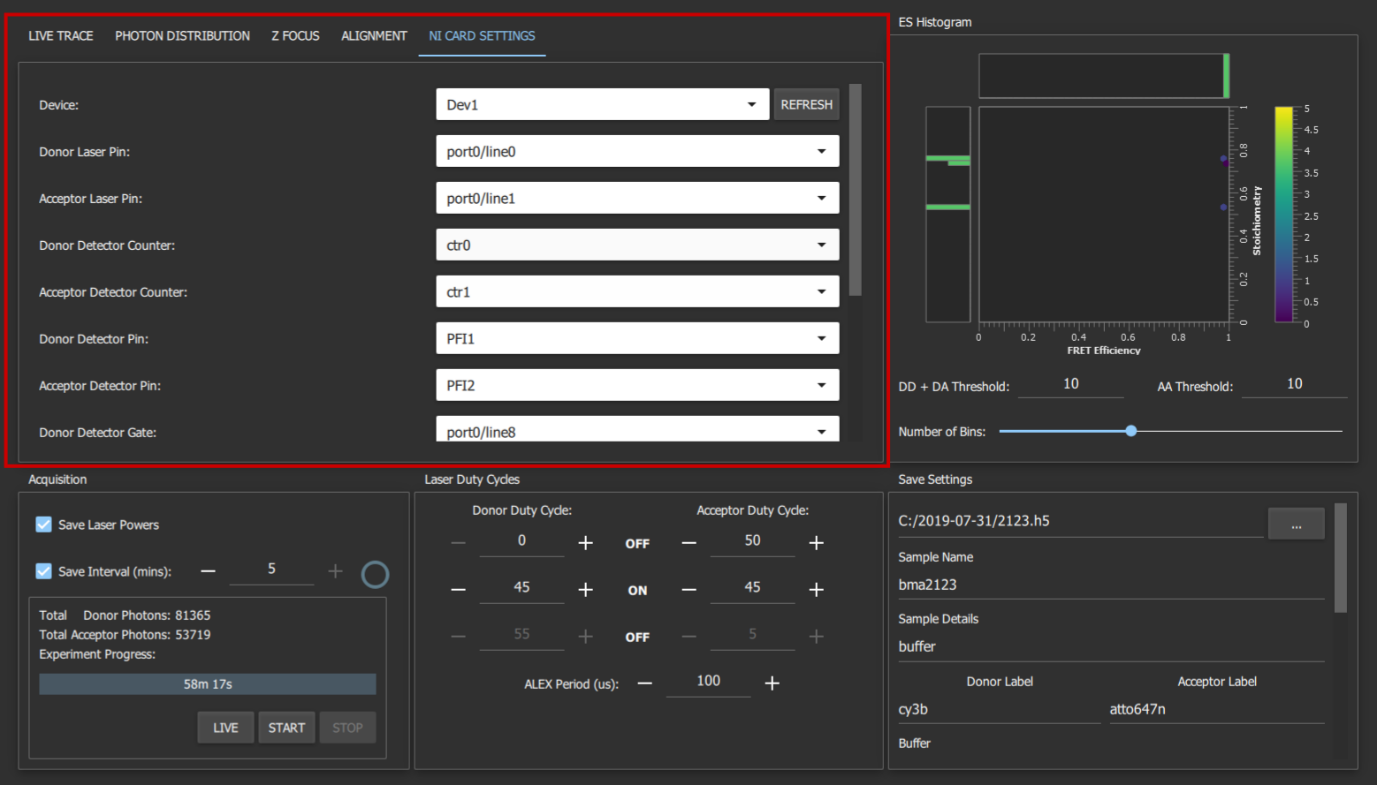

**Supplementary Fig. 16:** The NI Card Settings tab contains the advanced system settings that should not need to be changed unless you have constructed your system differently to our instructions. These settings are remembered between restarts.

The typical procedure for taking data with smOTTER is as follows; immersion oil and a coverslip is placed on the objective of the smfBox, a sample (10-100 μl) is applied and the lid is closed. The “z-focus” tab is used to position the z-stage such that the confocal volume is in the sample (see Supplementary Figure 13 for detail). Note that on slower computers the live trace may be slowed with the camera on, so the camera is stopped before acquisition. Then in the “live trace” tab, pressing the “live” button in the bottom left will begin displaying live data without saving. This is used to determine if the sample is at an appropriate concentration (1~ burst per second). If the sample is to be changed before acquisition, “stop” is pressed before opening the lid of the smfBox. When ready to acquire and save data, the experiment length and save frequency is entered, along with all save settings. On pressing “start”, smOTTER will begin to save acquired data.

***Supplementary Methods 23. Operational Software - LABVIEW***

The LABVIEW implementation has three separate VI’s for alignment, z-focussing, and acquisition, whereas smOTTER is capable of all and can save directly to HDF5 for quick analysis.

The prerequirements for the LABVIEW VI’s are:

[LabVIEW runtime 2016 64 bit](http://www.ni.com/download/labview-run-time-engine-2016/6067/en/) 
[VISA runtime engine 16.0 64 bit](http://www.ni.com/download/ni-visa-run-time-engine-16.0/6188/en/) 
[NI-DAQmx 16.0 64 bit](http://www.ni.com/en-gb/support/downloads/drivers/download.ni-daqmx.html#288254)

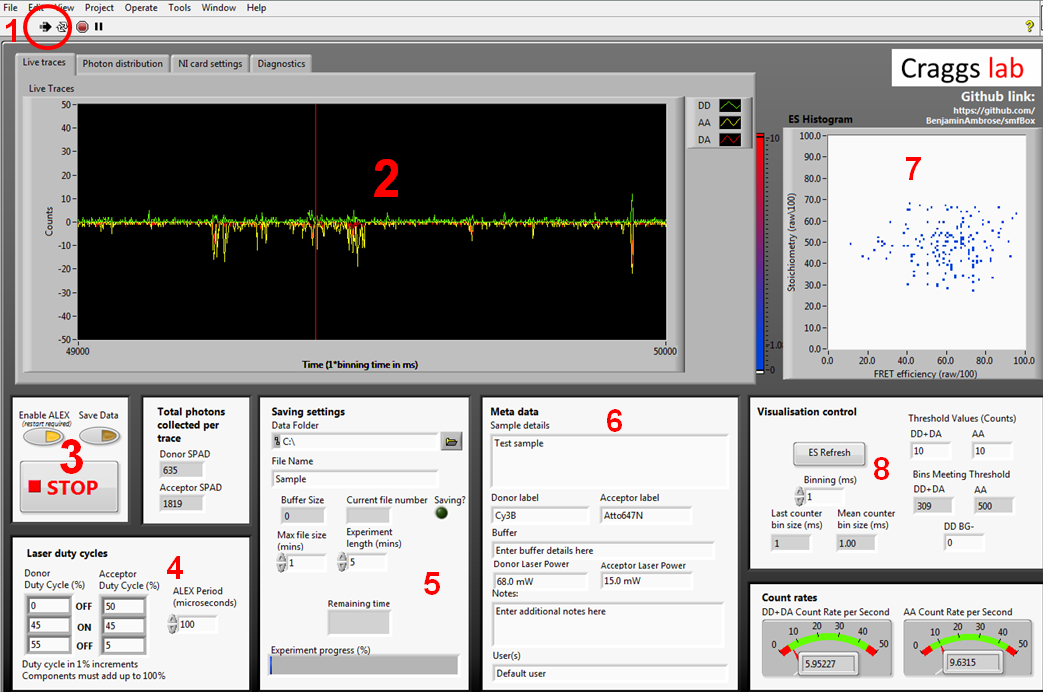

**Supplementary Fig. 17:** Front panel of the acquisition software. **1**. LABVIEW start button which begins ALEX and visualisation. **2.** Live time trace. Photons arriving at APD0 (donor) shown in green, and for ADP1 (acceptor) shown in red for photons under donor excitation and yellow for photons under acceptor excitation. **3**. Controls for ALEX, saving, and stopping. When you are ready to begin saving an acquisition, press the save data toggle during visualisation. **4.** ALEX controls. ALEX period controls the length of one full green-red alternation. The duty cycle boxes describe the amount of time each laser is off, then on, then off again for. This information is written into the metadata file, and pulled back out during file conversion to assign photons to channels. **5.** Location and name of saved files. You can set a max file size smaller than the experiment length and it will save as multiple files. **6.** Metadata typed here will be written into the meta data file, it will not affect the nature of the data itself. **7.** The visualisation control boxes affect the live 2D ES histogram on the bottom right. This histogram is built crudely and should not be used for rigorous data analysis, as it simply plots millisecond time bins exceeding the stated thresholds, rather than performing a full burst search. **8.** Controls for the live data visualisation. These do not affect the final saved data, only the live display.

The acquisition software controls the lasers and APDs and gives a live view of incoming smFRET data as well as creating the files needed for more in-depth analysis later. The first time you run the acquisition software you will need to go to the NI card settings tab and use the drop-down menus to select your NI card information, and the com ports each laser and APD are connected to.

Under normal operation; a user will typically place a sample on the scope, then press play (1) to check the quality/concentration before acquisition, we usually aim for 1-3 bursts per second to avoid co-incident events. If the sample needs to be changed again, press the stop button (3) before opening the lid of the smfBox to prevent ambient light from flooding the APD's. When happy with the sample, set the acquisition length and save location in (5), add metadata in (6), then press "Save Data" to begin the acquisition. 
The diagnostics tab can be used to determine any problems with the software or hardware, and the photon distribution tab will give a live histogram of when photons have been detected with respect to the ALEX period. This can be a useful diagnostic of whether the lasers are being controlled properly without having to create a file for analysis first.

***Supplementary Methods 4. Data Files***

The LABVIEW acquisition software saves raw txt files which contain the timestamp of each photon in one column and the detector it arrived at in a second column. For usage in analysis we have created python scripts which can convert these raw data to the open source photon-HDF5 format^15^. This format contains not only the raw data but also can be written with sufficient metadata that the reader knows i. the origin and nature of the sample and experiment the data was acquired from and ii. Information required to interpret it. In this case the data originates from two unpolarised detectors acquired with a μs-ALEX scheme, and the script will write an HDF5 file which contains information for any analysis software to interpret it as such.

The scripts provided can convert a single TXT to a single HDF5, multiple TXT’s to multiple HDF5’s (batch conversion), or multiple TXT’s for single HDF5’s in the case of repeat acquisitions which you wish to analyse as a single dataset.

These conversion scripts are available as Jupyter Notebooks, as the analysis we provide is with the FRETBursts python module in Jupyter. However there is no reason why the code should not work in any other python environment as long as it has the phconvert module.

The standalone C++ software smOTTER will save directly to an HDF5.

***Supplementary Methods 5. DNA sequences***

| **Name** | **Sequence** |
| --- | --- |
| 1a | 5’- GAG CTG AAA GTG TCG AGT TTG TTT GAG TGT **T**TG TCT GG - 3’  3’- CTC GAC T**T**T CAC AGC TCA AAC AAA CTC ACA AAC AGA CC - 5’ - biotin |
| 1b | 5’- GAG CTG AAA GTG TCG AGT TTG T**T**T GAG TGT TTG TCT GG - 3’  3’- CTC GAC T**T**T CAC AGC TCA AAC AAA CTC ACA AAC AGA CC - 5’ - biotin |
| 1c | 5’- GAG CTG AAA GTG TCG AGT **T**TG TTT GAG TGT TTG TCT GG - 3’  3’- CTC GAC T**T**T CAC AGC TCA AAC AAA CTC ACA AAC AGA CC - 5’ - biotin |

**Supplementary Table 5:** Accurate FRET standard sequences from previous study^5^. Highlighted T’s in green and red represent C2 amino modified thymine residues labelled with Atto-550 and Atto-647N respectively. Acceptor strands are labelled with 5’ biotins for surface immobilisation in the case of TIRF measurements in the original study (not used here).

| **Name** | **Sequence** |
| --- | --- |
| MJW1 | 5’ -T(C6 amino)-GGATTAAAAAAAAAAAAAAAAAAAAAAAAAAAAAAAAATCCAAAGGATGTATGGTAATGGGACGA  AGAATGAGG 3’ |
| MJW2 | 5’ CCTCATTCTTCGTCCCATTACCA-T(C6 amino)-ACATCC 3’ |
| MJW3 | 5’ CCTCATTCTTCGTCCCAT-T(C6 amino)-ACCATACATCC 3’ |
| MJW4 | 5’ CCTCATTCTTCG-T(C6 amino)-CCCATTACCATACATCC 3’ |

**Supplementary Table 6:** DNA Hairpin Sequences. MJW1 was labelled with Cy3B, and MJW2, MJW3, MJW4 were labelled with Atto647N, then annealed to produce the high, mid, and low FRET hairpins respectively.

***Supplementary Methods 6. Accurate FRET validation***

Double stranded DNA’s labelled with Atto550 and Atto647n 23, 15, and 11-bp apart as described in Hellenkamp et al., (2018) were diluted to approximately 0.1 nM in a buffer consisting of 5 mM Tris, pH 7.5, 20 mM MgCl_2_ and 5 mM NaCl. 20 μl of diluted sample was placed on the smfBox on a coverslip passivated with BSA and covered with the cap of a micro centrifuge tube to prevent evaporation.

515 nm and 635 nm lasers were alternated with a 45/5/45/5 μs green/off/red/off cycle for two, half-hour acquisitions per sample. Time stamped data from the two APD’s were converted to the open-source photon-HDF5 file format using custom Jupyter Notebooks (provided in supplementary data) with the phconvert python module.

Analysis was done using Jupyter Notebooks using the FRETBursts^16^ python module (0.6.5), and Jupyter Notebooks (available in the supplemental material) following the method described in Hellenkamp et al., (2018). First “Correction Factor Finder Alpha-Delta.ipynb” is used followed by “Correction Factor Finder Gamma-Beta.ipynb” to determine correction factors, then “FRET Analysis.ipynb” is used to find FRET efficiencies of each sample. A brief description of the overall process is as follows: Background in each channel was estimated by means of an exponential fit of inter-photon delays greater than 1.5 ms, calculated for every 5 minutes of data. Bursts were identified using either an all photon sliding window algorithm, or dual channel burst search (DCBS) algorithm previously described^24^ with L=10 and F=45 for both channels, and background was subtracted. For the spectral cross talk factors, data from all experiments were combined and an all photon search was used, and only bursts with >50 photons in any channel were selected, and ES plots were made (Supplementary Equations 1 and 2). Bursts with a stoichiometry >0.95 were selected as the donor only population and <0.175 for the acceptor only population. α and δ factors were calculated from E and S single Gaussian fits of donor only and acceptor only respectively using Supplementary Equations 3 and 4. A dual channel burst search (DCBS) was then used to extract doubly labelled bursts from each 30-minute acquisition, and bursts with >50 photons under green excitation were used to find E and S with single Gaussian fits. 2D Gaussian positions of each 30-minute data set were then plotted together and fitted with Supplementary Equation 5 to obtain γ and β. 30-minute data sets were then combined for each sample, bursts extracted using DCBS with accurate FRET parameters applied, selected for >50 photons under green excitation, and FRET efficiency obtained via a single Gaussian fit.

***Supplementary Results 1. Accurate FRET Results***

|  | Hellenkamp et al., 2018 | smfBox Data | | |
| --- | --- | --- | --- | --- |
| Sample | E | E | σ | N bursts |
| 1a | 0.15 ±0.02 | 0.17 | 0.07 | 694 |
| 1b | 0.56 ±0.03 | 0.57 | 0.10 | 717 |
| 1c | 0.76 ±0.015 | 0.77 | 0.07 | 799 |

**Supplementary Table 7:** Summary statistics for the accurate FRET validation. The means and standard deviations of FRET efficiencies from all participants in the benchmarking study are shown, alongside the FRET efficiency mean (E) and width (σ) of a Gaussian fit to data from the smfBox.

***Supplementary Note 1. Asymmetric ALEX***

The rate and duty cycle of the green and red lasers of the smfBox can be completely customised using either acquisition software. It is even possible to leave one laser on all the time (for PAX, for example), or to leave one laser permanently off if ALEX is not required. For the experiments in this study we chose a duty cycle length of 100 μs, as this is fast enough to allow several observations with either laser per molecule (observation time of ~1 ms). If there are too few alternations then it reduces the ability of algorithms like the dual channel burst search to discriminate between molecules which have bleached during observation. However, the lasers have a rise/fall time of several microseconds (Supplementary Fig. 16), so repeating too fast would decrease the total time the lasers are on at full power. Furthermore, in either the case of converting to HDF5 from the LABVIEW software, or in directly saving to HDF5 in smOTTER, the details of the ALEX cycle are contained within the metadata, which is then read out by FRETBursts^16^, so analysis is not further complicated by changing any of these parameters.

We have demonstrated the capabilities and effects of altering the ALEX cycle with a number of asymmetric excitation schemes, in which the amount of time either laser is on for is not equal. Typically ALEX schemes are symmetrical, however there is not strictly any reason why the red laser need be on for the same amount of time as the green. In fact, we argued that since all the information for the FRET efficiency comes under green excitation, it makes sense to maximise the time this laser is on. The four datasets shown (Supplementary Fig. 18) were taken using duplex 1b as described in the main text. A dual channel burst search (F = 15, m = 10) was used for analysis, and a burst selection of 50 photons under green excitation and 50 photons under red excitation.

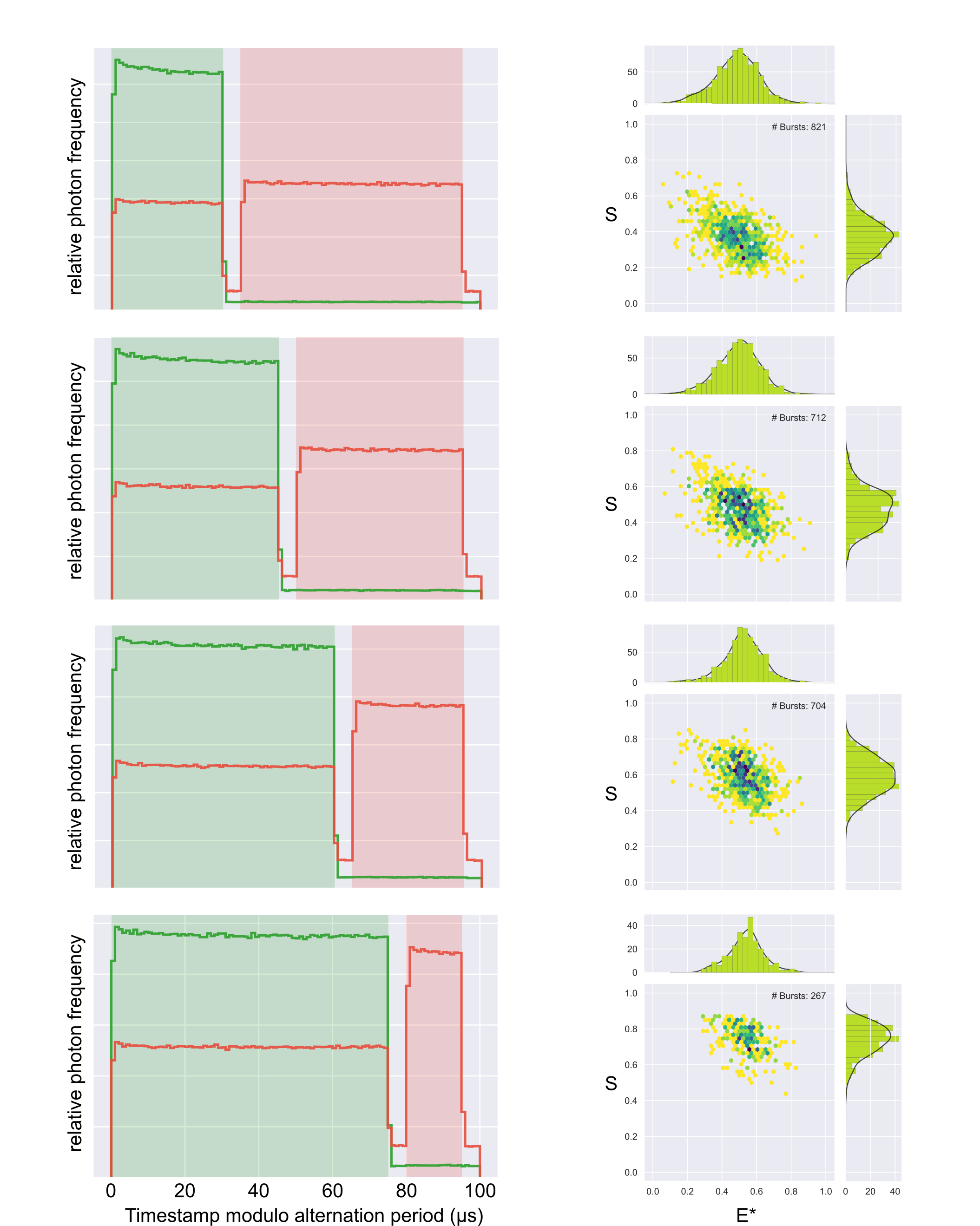

**Supplementary Fig. 18:** Four different ALEX schemes. On the left are photon arrival time plots with detected photons as green and red lines and shaded boxes representing which laser was on. On the right are uncorrected ES Histograms of DNA standard 1b measured with these ALEX schemes. From top to bottom the green/red excitation times are 30/60 μs, 45/45 μs, 60/30 μs, and 75/15 μs. In all cases a 5 μs wait time was left between each laser.

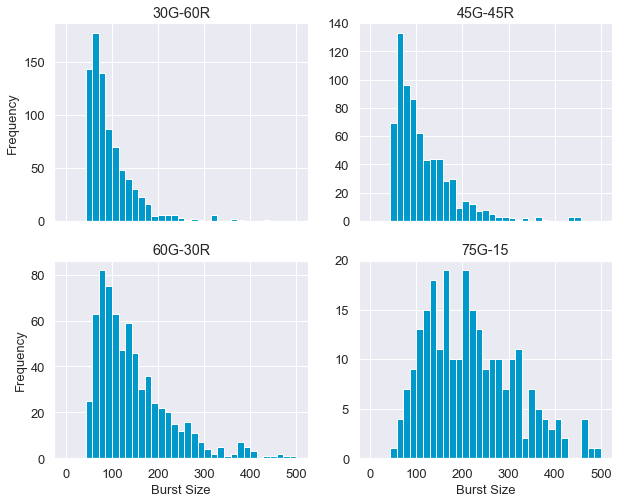

**Supplementary Fig. 19:** Burst size histograms for each excitation scheme. Plotted are the number of photons under green excitation after the selection of >50 photons, which is why there are no bursts <50 photons shown.

Changing the relative amounts of excitation by either laser shows a predictable trend (Supplementary Figure 18). The S position in each ES plot increases with the length of green excitation, due to the increased number of photons detected under green excitation. However, two other effects become apparent. Firstly, the width of the distribution in E decreases due to the decreased contribution of shot-noise from the increased number of photons acquired. Secondly, the number of doubly labelled bursts which are found by the burst search algorithm decreases due to the reduced information under acceptor excitation, which is required to confidently assert that any burst is doubly labelled. From these two effects it is easy to see that there may be an optimum excitation scheme, which may not necessarily be symmetrical, since the quantity of information obtained under donor excitation is greater than that obtained under acceptor excitation. As expected, we can see that the burst size histogram shifts to increasing number of photons under green excitation when the green excitation period is increased (Supplementary Figure 19), consistent with the reduction in width in FRET efficiency histogram due to decreased shot noise.

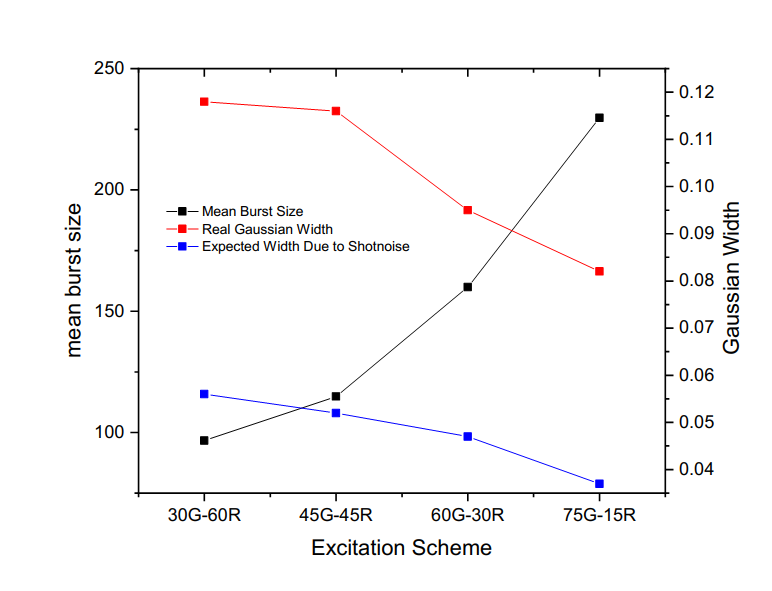
**Supplementary Fig. 20.** Burst statistics under each excitation scheme, showing: the mean number of photons under green excitation (black); the width of the Gaussian fitted to the 1D FRET efficiency distribution (red); and the expected Gaussian width computed from the burst size distribution (blue).

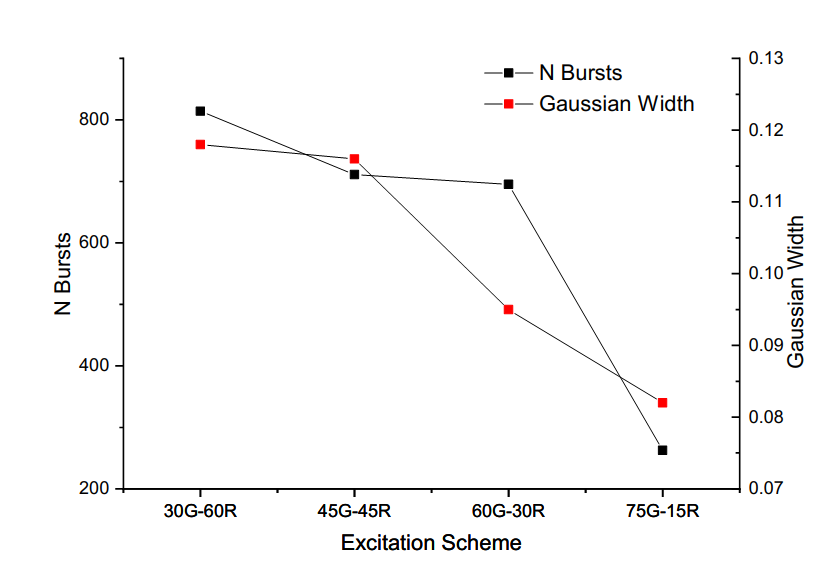
**Supplementary Fig. 21:** Four different excitation schemes with the number of bursts found by a DCBS and the width of the Gaussian distributions fitted to the FRET efficiencies.

To further emphasise this point, we see that Gaussian width decreases with increasing burst size, as can be predicted by a binomial computation of shot noise from the burst size distribution (Supplementary Figure 20). Note that the width of the Gaussian distribution fitted to the experimental data is larger than predicted from shot noise alone, which is likely due to background contribution, and flexing of the DNA molecule. This discrepancy has been previously discussed^4,24,25^ . Finally in Supplementary Figure 21 we can see the effects of each excitation scheme on both Gaussian width and number of detected bursts. Whilst more testing may be necessary to fully explore this relationship, it can be seen phenomenologically that increasing donor excitation to twice that of the acceptor can markedly decrease the width of the obtained FRET distribution without having to resort to increasing laser power (and hence increasing photobleaching), whilst having little to no effect on the ability to confidently assert that a burst is doubly labelled. From this plot we would recommend a ‘sweet-spot’ of 60G-30R for the duty cycle, to both maximise the number of detected bursts and minimise the shot noise.

***Supplementary Note 2. Verifying Dynamics***

Typically, the PDA method is supplemented with a burst-wise test for dynamics which can verify the hypothesis that bursts of intermediate FRET efficiency originate from molecules undergoing conformational changes, as PDA does not directly detect dynamics, but rather uses them to compute kinetic rates. If fluorescence life-times are available then an E-tau plot can discriminate dynamic and static bursts via deviation from a “static FRET” line^26^. If only intensities are available, then either the burst variance analysis (BVA) method^27^ or two channel kernel density estimator^25^ (2CDE) can be used to verify dynamics using photon arrival time statistics. Here we show the FRET-2CDE method as implemented using PAM^18^ applied to the hairpins in this study, using the static dsDNA’s as a control to demonstrate not only supporting evidence that the hairpins are dynamic, but that this form of analysis is also possible with the smfBox platform.

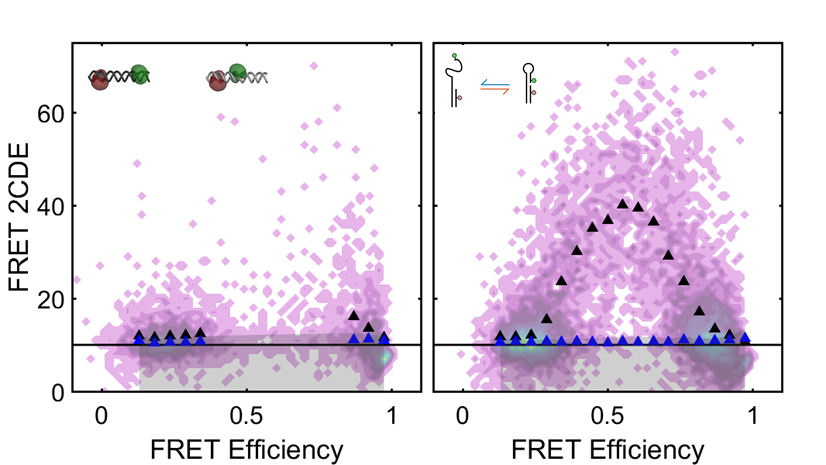

**Supplementary Fig. 22:** 2CDE plots of a static dsDNA mixture (left) and a dynamic hairpin (right). FRET-2CDE analysis^28^ can be done in PAM on data from the smfBox in order to verify the presence of dynamics in the sample. A high 2CDE value (average shown in black triangles) suggests that bursts are dynamic, ie. Interconverting between FRET efficiencies during observation, whereas low 2CDE bursts along a straight line (blue triangles) suggest a static FRET efficiency.

***Supplementary Note 3. Possible Expansions***

Due to the modular nature of the smfBox, it can be readily expanded for use in other confocal based techniques beyond the scope of this paper. Various possible additions include:

2 axis scanning stage – eg from Piezoconcept / MadCity Labs

Pulsed lasers and controllers – eg from Picoquant

Time correlated single-photon counting electronics (picosecond resolution) – eg from Picoquant

Additional APDs (Excelitas) and polarisation optics (eg Thorlabs).

***Supplementary Equations***

$Proximity Ratio, PR= E* = \frac{DA}{DD+DA}$	(1)

$S= \frac{DD+DA}{DD+DA+AA}$	(2)

$\alpha= \frac{E_{app}^{Donly}}{1- E_{app}^{Donly}}$	(3)

$\delta= \frac{S_{app}^{Aonly}}{1- S_{app}^{Aonly}}$	(4)

$S_{app}^{DA}=\frac{1}{\left( 1+ \gamma\beta+(1-\gamma)\beta E_{app}^{DA} \right)}$	(5)

***Supplementary References***

25.	Antonik, M., Felekyan, S., Gaiduk, A. & Seidel, C. A. M. Separating Structural Heterogeneities from Stochastic Variations in Fluorescence Resonance Energy Transfer Distributions via Photon Distribution Analysis. *J. Phys. Chem. B* **110**, 6970–6978 (2006).

26.	Kalinin, S., Valeri, A., Antonik, M., Felekyan, S. & Seidel, C. A. M. Detection of Structural Dynamics by FRET: A Photon Distribution and Fluorescence Lifetime Analysis of Systems with Multiple States. *J. Phys. Chem. B* **114**, 7983–7995 (2010).

27.	Torella, J. P., Holden, S. J., Santoso, Y., Hohlbein, J. & Kapanidis, A. N. Identifying molecular dynamics in single-molecule FRET experiments with burst variance analysis. *Biophys. J.* **100**, 1568–1577 (2011).

28.	Tomov, T. E. *et al.* Disentangling Subpopulations in Single-Molecule FRET and ALEX Experiments with Photon Distribution Analysis. *Biophys. J.* **102**, 1163–1173 (2012).
