## Supplementary material for "The smfBox: an open-source platform for single-molecule FRET": Jupyter Notebooks and data for accurate FRET: 1a - FRET Analysis 1.3.ipynb

```
# %% [markdown]
# smFRET Analysis

This notebook is for simple analysis of smFRET data, starting with an hdf5 file and ending with a FRET efficiency histogram that can be fitted with a gaussians. Burst data can be exported as a .csv for analysis elsewhere.

You can analysis uncorrected data if you are simply looking for relative changes in the conformational ensemble, or accurate FRET correction parameters can be supplied if you want FRET efficiencies that can be converted to distances.

# %% [markdown]
# Import packages

# %% [code]
from fretbursts import *
sns = init_notebook()
import lmfit
import phconvert
import os
import matplotlib.pyplot as plt
import matplotlib as mpl
from fretbursts.burstlib_ext import burst_search_and_gate

# %% [markdown]
# Name and Load in data

First name the data file and check it exists, it will look for the file starting from wherever this notebook is saved.

# %% [code]
filename = "definitiveset/1ax.hdf5"
if os.path.isfile(filename):
    print("File found")
else:
    print("File not found, check file name is correct")

# %% [markdown]
Load in the file and set correction factors. If you aren't using accurate correction factors, set d.leakage and d.dir_ex to 0 and d.gamma to 1

You may get warnings that some parameters are not defined in the file, this is fine as they will be defined in this workbook anyway

# %% [code]
d = loader.photon_hdf5(filename)
for i in range(0, len(d.ph_times_t)): #sorting code
    indices = d.ph_times_t[i].argsort()
    d.ph_times_t[i], d.det_t[i] = d.ph_times_t[i][indices], d.det_t[i][indices]
d.leakage = 0.081 #alpha
d.dir_ex = 0.076 #delta
d.gamma = 0.856
d.beta = 0.848
#d.leakage = 0.
#d.dir_ex = 0.
#d.gamma = 1.

# %% [markdown]
# Check alternation cycle is correct

We need to check that the ALEX parameters defined in the HDF5 file are appropriate for the laser cycle used in the experiment. If this is correct, the following histogram should look correct. It is a combined plot of every photon that arrives over the supplied alternation periods. 

# %% [code]
bpl.plot_alternation_hist(d)

# %% [markdown]
IF THE ABOVE HISTOGRAM LOOKS CORRECT: then run loader.alex_apply_period, which rewrites the time stamps into groups based on their excitation period. If you want to change the alternation period after this you will have to reload the data into FRET bursts.

IF THE ABOVE HISTOGRAM LOOKS WRONG: then the supplied alternation parameters do not match up to the alternation of the lasers in the data. This could be because the lasers were actually on a different alternation, or because the data set doesn't start at zero so is frame shifted etc.

In this case, you can un-hash the code below and alter the parameters manually.

# %% [code]
#d.add(det_donor_accept = (0, 1),
#    alex_period = 10000,
#    offset = 0,
#    D_ON = (0, 4500),
#    A_ON = (5000, 9500))

# %% [code]
loader.alex_apply_period(d)
time = d.time_max
print('Total data time = %s'%time)

# %% [markdown]
The following will plot a time trace of the first second of your experiment.

# %% [code]
dplot(d, timetrace, binwidth=1e-3, tmin=0, tmax=15, figsize=(8,5))
plt.xlim(0,1);
plt.ylim(-45,45);
plt.ylabel(" Photons/ms in APD0       Photons/ms in APD1");

# %% [markdown]
# Background Estimation

# %% [markdown]
Background estimation works by plotting log of photons by the delay between them, assuming a poisson distribution of photon arrivals and fitting a line. The plot will contain single molecule bursts however, so a threshold (in microseconds) has to be defined where the fit begins. 

The variable "time_s" defines the size of the windows in which background is recalculated. Lower values will make the experiment more accurately sensisitive to fluctuations in background however higher values will give more photons with which to calculate a more precise average background. If the fit fails you may need to increase this value.

# %% [code]
threshold = 1500
d.calc_bg(bg.exp_fit, time_s=300, tail_min_us=(threshold),)
dplot(d, hist_bg, show_fit=True)

# %% [markdown]
This code will plot the calculated background in each window and acts as a good reporter of whether anything major has happened to the solution over the time course of the experiment

# %% [code]
dplot(d, timetrace_bg);

# %% [markdown]
# Burst Searching and Selecting

"d.burst_search()" can be used to do an all photon burst search (APBS), however "burst_search_and_gate(d)" will apply a DCBS / dual channel burst search (Nir 2006), this effectively does independent searches in the DD+DA channel and the AA channel, and then returns the intersecton of these bursts, ensuring that any FRET information is only included whilst an acceptor is still active in the detection volume.

The two numbers given to "F=" are the signal to background threshold in the DD+DA and AA channels respectively. If your background is particularly high in one but not the other you may want to change these independently.

# %% [code]
bursts=burst_search_and_gate(d, F=(20, 20), m=10, mute=True)

# %% [markdown]
The following will plot a graph of burst number vs sizes which can inform your selection thresholding. 

# %% [code]
sizes = bursts.burst_sizes_ich(add_naa=True)
plt.hist(sizes, cumulative=-1, bins = 30, range = (0, 100), histtype="stepfilled", density=False)
plt.xlabel('Burst size (n photons)')
plt.ylabel("N bursts with > n")

# %% [markdown]
We can now set thresholds on how many photons we want in each burst, this can be done on all channels together, or just one channel. Thresholding DD+DA will reduce the width in E, thresholding in AA will ensure there are no donor only bursts.

# %% [code]
bursts = bursts.select_bursts(select_bursts.size, add_naa=True, th1=20,) #all channels
bursts = bursts.select_bursts(select_bursts.size, th1=50) #DD + DA
bursts = bursts.select_bursts(select_bursts.naa , th1=10) #AA

# %% [markdown]
# Histograms

Now we can start plotting and fitting the data

# %% [code]
g=alex_jointplot(bursts)

# %% [markdown]
This will fit a gaussian to the E values.

If you set pdf=True then the data will be displayed as a probability density function, pdf=False will give it as bursts instead

# %% [code]
model = mfit.factory_gaussian()
model.set_param_hint('center', value=1.1, vary=True)
bursts.E_fitter.fit_histogram(model=model, verbose=False, pdf=False)
dplot(bursts, hist_fret, pdf=False, show_model=True, show_fit_value=True, fit_from='center');
bursts.E_fitter.params

# %% [markdown]
This will export the data to a .csv file, type the save location between the ""'s. This csv file can be opened in excel or origin and contains information about each burst, most importantly E and S but also things like burst length and width.

# %% [code]
csvfile = "1ax.csv"
burstmatrix = bext.burst_data(bursts)
burstmatrix.to_csv(csvfile)
```
