## Supplementary material for "The smfBox: an open-source platform for single-molecule FRET": Jupyter Notebooks and data for accurate FRET: Converter - Batch txt to Photon-HDF5 3.2 (Multiple Runs).ipynb

```
# %% [markdown]
# Multiple run converter

This notebook will convert a whole folder of files each to their respective HDF5's, incorporating metadata it finds. It works off the existing file structure created by labview saving. Do not move any files outside of the folder or it will not work, see the website for more details.

# %% [code]
import phconvert as phc
import csv
phc.__version__
import numpy as np
import os

# %% [markdown]
Point this at a folder which contains folders, which contain data and metadata files together. It will find the data files and metadata files in each folder and create a single HDF5 for each, named according the name you gave the data file (it will automatically chop off the annoying numbers at the beginning and end).

# %% [code]
parentfolder = "accurateFRETstandards/" #THE DASH AT THE END IS IMPORTANT

# %% [code]
def meta(number):
    param = metadata[metadata.find(params[number])+len(params[number]):metadata.rfind(params[number+1])]
    return(param)
for folder in next(os.walk(parentfolder))[1]:
    filenames = []
    for file in os.listdir(parentfolder+folder):        
        if file.endswith("T.txt"):
            filenames.append(file)
        if file.endswith("etadata.txt"):
            metafile = os.path.join(parentfolder+folder+"/"+file)
        else:
            continue
    savename = os.path.join(parentfolder+filenames[0][:-8][4:]+".hdf5")
    detectors = np.empty([0], dtype=int)
    timestamps = np.empty([0], dtype=int)
    print("Converting " + filenames[0][:-8][4:])
    for x in filenames:
        file = os.path.join(parentfolder+folder+"/"+x)
        with open(file) as inf:
            reader = csv.reader(inf, delimiter="	")
            ftimestamps = list(zip(*reader))[0]    
        with open(file) as inf:
            reader = csv.reader(inf, delimiter="	")
            fdetectors = list(zip(*reader))[1]
        fdetectors = np.asarray(fdetectors)
        ftimestamps = np.asarray(ftimestamps)
        ftimestamps = np.int64(ftimestamps)
        fdetectors = np.uint8(fdetectors)
        timestamps = np.concatenate([timestamps, ftimestamps])
        detectors = np.concatenate([detectors, fdetectors]) 
    timestamps = np.int64(timestamps)
    detectors = np.uint8(detectors)
    with open(metafile, 'r') as file:
        metadata = "".join(line.rstrip() for line in file)
    params = ('Acquistion time per file (mins): ', 'Donor Laser Intensity: ', 'Acceptor Laser Intensity: ', 'Users: ', 'Approximate Fluorophore Concentration: ',
             'Sample details: ', 'Donor Label: ', 'Acceptor Label: ', 'Buffer: ', 'Notes: ',
            'D_ON: ', 'D_OFF: ', 'A_ON: ', 'A_OFF: ',)
    D_ON, D_OFF, A_ON,= int(meta(10)), int(meta(11)), int(meta(12))
    A_OFF = int(metadata[metadata.find(params[13])+len(params[13]):])
    sample_name = meta(5)
    buffer_name = meta(8)
    dye_names = meta(6) + "," + meta(7)
    timestamps_unit = 10e-9  # 10 ns, units are always S.I.
    photon_data = dict(
        timestamps=timestamps,
        detectors=detectors,
        timestamps_specs={'timestamps_unit': timestamps_unit})
    setup = dict(
        num_pixels = 2,                   # using 2 detectors
        num_spots = 1,                    # a single confoca excitation
        num_spectral_ch = 2,              # donor and acceptor detection 
        num_polarization_ch = 1,          # no polarization selection 
        num_split_ch = 1,                 # no beam splitter
        modulated_excitation = False,     # CW excitation, no modulation 
        excitation_alternated = [True],  # CW excitation, no modulation 
        lifetime = False,                 # no TCSPC in detection    
        excitation_wavelengths = [515e-9, 635e-9],         # List of excitation wavelenghts
        excitation_cw = [True],                    # List of booleans, True if wavelength is CW
        detection_wavelengths = [580e-9, 690e-9],  # Nominal center wavelength 
    )
    identity = dict(
        author= meta(3),
        author_affiliation="University of Sheffield") #CHANGE THIS WHERE NECESSARY
    measurement_specs = dict(
        measurement_type = 'smFRET-usALEX',
        detectors_specs = {'spectral_ch1': [0],  # list of donor's detector IDs
                           'spectral_ch2': [1]},  # list of acceptor's detector IDs,
        alex_period = 10000,
        alex_offset = 0,
        alex_excitation_period1 = (D_ON, D_OFF),
        alex_excitation_period2 = (A_ON, A_OFF)
        )
    photon_data['measurement_specs'] = measurement_specs
    data = dict(
        description=meta(9),
        photon_data = photon_data,
        setup=setup,
        identity=identity,
    )
    phc.hdf5.save_photon_hdf5(data, h5_fname=savename, overwrite=True)
print("all done")

# %% [code]
```
