## Supplementary material for "The smfBox: an open-source platform for single-molecule FRET": Jupyter Notebooks and data for accurate FRET: Converter - Multiple TXT to Photon-HDF5 3.2 (Single Run, multiple files).ipynb

```
# %% [markdown]
# Multiple file run converter

This notebook will find multiple files from a single run within a folder, and the metadata file then convert them. This is useful if you have set the acquisition software to take for example, an hour long acquisition, but with 10 minute long files.

It works directly off of the naming and file structure created by the acquisition software. Do not move any files outside of the folder or it will not work, see the website for more details.

# %% [code]
import phconvert as phc
import csv
phc.__version__
import numpy as np
import os

# %% [markdown]
Start by naming the folder your files are in, this code will then find all of the files in it, and report which it has found.

# %% [code]
folder = 'accuratefretstandards/' #put the RAW filename here
savename = 'accratefretstandards/1a.hdf5' #put the save location / name herefilenames


filenames = [] #LEAVE THIS EMPTY!!!
for file in os.listdir(folder):        
    if file.endswith("T.txt"):
        datafile = file
        filenames.append(folder+file)
    if file.endswith("etadata.txt"):
        metafile = (os.path.join(folder, file))
    else:
        continue
print("Found data files:")
print(filenames)
print("Found metadata:")
print(metafile)

# %% [markdown]
Read in the columns from each file and concatenate them. It will report the first and last photons from every file read in. If they do not logically follow on from each other then something has gone wrong.

# %% [code]
detectors = np.empty([0], dtype=int)
timestamps = np.empty([0], dtype=int)
for file in filenames:
    with open(file) as inf:
        reader = csv.reader(inf, delimiter="	")
        ftimestamps = list(zip(*reader))[0]    
    with open(file) as inf:
        reader = csv.reader(inf, delimiter="	")
        fdetectors = list(zip(*reader))[1]
    fdetectors = np.asarray(fdetectors)
    ftimestamps = np.asarray(ftimestamps)
    ftimestamps = np.int64(ftimestamps)
    fdetectors = np.uint8(fdetectors)
    print(ftimestamps)
    timestamps = np.concatenate([timestamps, ftimestamps])
    detectors = np.concatenate([detectors, fdetectors]) 
timestamps = np.int64(timestamps)
detectors = np.uint8(detectors)
print("done")

# %% [markdown]
Now we can create an HDF5 file by combining the data with the metadata

# %% [code]
with open(metafile, 'r') as file:
  metadata = "".join(line.rstrip() for line in file)
params = ('Acquistion time per file (mins): ', 'Donor Laser Intensity: ', 'Acceptor Laser Intensity: ', 'Users: ', 'Approximate Fluorophore Concentration: ',
             'Sample details: ', 'Donor Label: ', 'Acceptor Label: ', 'Buffer: ', 'Notes: ',
            'D_ON: ', 'D_OFF: ', 'A_ON: ', 'A_OFF: ',)
def meta(number):
    param = metadata[metadata.find(params[number])+len(params[number]):metadata.rfind(params[number+1])]
    return(param)
D_ON, D_OFF, A_ON,= int(meta(10)), int(meta(11)), int(meta(12))
A_OFF = int(metadata[metadata.find(params[13])+len(params[13]):])
sample_name = meta(5)
buffer_name = meta(8)
dye_names = meta(6) + "," + meta(7)
timestamps_unit = 10e-9  # 10 ns, units are always S.I.
photon_data = dict(
    timestamps=timestamps,
    detectors=detectors,
    timestamps_specs={'timestamps_unit': timestamps_unit})
setup = dict(
    num_pixels = 2,                   # using 2 detectors
    num_spots = 1,                    # a single confoca excitation
    num_spectral_ch = 2,              # donor and acceptor detection 
    num_polarization_ch = 1,          # no polarization selection 
    num_split_ch = 1,                 # no beam splitter
    modulated_excitation = False,     # CW excitation, no modulation 
    excitation_alternated = [True],  # CW excitation, no modulation 
    lifetime = False,                 # no TCSPC in detection    
    excitation_wavelengths = [515e-9, 635e-9],         # List of excitation wavelenghts
    excitation_cw = [True],                    # List of booleans, True if wavelength is CW
    detection_wavelengths = [580e-9, 690e-9],  # Nominal center wavelength 
)
identity = dict(
    author= meta(3),
    author_affiliation="University of Sheffield") #CHANGE THIS WHERE NECESSARY
measurement_specs = dict(
    measurement_type = 'smFRET-usALEX',
    detectors_specs = {'spectral_ch1': [0],  # list of donor's detector IDs
                       'spectral_ch2': [1]},  # list of acceptor's detector IDs,
    alex_period = 10000,
    alex_offset = 0,
    alex_excitation_period1 = (D_ON, D_OFF),
    alex_excitation_period2 = (A_ON, A_OFF)
    )
photon_data['measurement_specs'] = measurement_specs
data = dict(
    description=meta(9),
    photon_data = photon_data,
    setup=setup,
    identity=identity,
)
phc.hdf5.save_photon_hdf5(data, h5_fname=savename, overwrite=True)

# %% [code]
```
