## Supplementary material for "The smfBox: an open-source platform for single-molecule FRET": Jupyter Notebooks and data for accurate FRET: Converter - Single TXT to Photon-HDF5 3.2 (Single Run, single file).ipynb

```
# %% [markdown]
# Converting raw .txt files from labview to fretbursts compatible HDF5 files

This notebook will convert the .txt files written by our labview acquisition software into HDF5 files which can be read into other FRETbursts notebooks. 

At the moment it generates dummy metadata but at some point I will get it to also read in the meta data files generated by labview and write those into the final HDF5 file.

# %% [code]
import phconvert as phc
import csv
phc.__version__
import numpy as np
import os

# %% [markdown]
Name the exact files containing the raw data and meta data and specify the save file location and name.

# %% [code]
raw = '0000/1a.txt'
metafile = "0000/metadata.txt"
savename = '0000/1a.hdf5'

# %% [markdown]
Check files are named properly

# %% [code]
try:
    with open(raw): pass
    print('Data file found, you can proceed.')
except IOError:
    print('ATTENTION: Data file not found, please check the filename.\n'
          '           (current value "%s")' % raw)
try:
    with open(metafile): pass
    print('Metadata file found, you can proceed.')
except IOError:
    print('ATTENTION: Metadata file not found, please check the filename.\n'
          '           (current value "%s")' % raw)

# %% [markdown]
Read in columns from the data files

# %% [code]
with open(raw) as inf:
    reader = csv.reader(inf, delimiter="	")
    timestamps = list(zip(*reader))[0]
with open(raw) as inf:
    reader = csv.reader(inf, delimiter="	")
    detectors = list(zip(*reader))[1]
detectors = np.asarray(detectors)
timestamps = np.asarray(timestamps)
timestamps = np.int64(timestamps)
detectors = np.uint8(detectors)

# %% [markdown]
Read in the metadata 

# %% [code]
with open(metafile, 'r') as file:
  metadata = "".join(line.rstrip() for line in file)
params = ('Acquistion time per file (mins): ', 'Donor Laser Intensity: ', 'Acceptor Laser Intensity: ', 'Users: ', 'Approximate Fluorophore Concentration: ',
             'Sample details: ', 'Donor Label: ', 'Acceptor Label: ', 'Buffer: ', 'Notes: ',
            'D_ON: ', 'D_OFF: ', 'A_ON: ', 'A_OFF: ',)
def meta(number):
    param = metadata[metadata.find(params[number])+len(params[number]):metadata.rfind(params[number+1])]
    return(param)

# %% [markdown]
Read in the info from the metadata and use it to build the HDF5

# %% [code]
D_ON, D_OFF, A_ON,= int(meta(10)), int(meta(11)), int(meta(12))
A_OFF = int(metadata[metadata.find(params[13])+len(params[13]):])
sample_name = meta(5)
buffer_name = meta(8)
dye_names = meta(6) + "," + meta(7)
timestamps_unit = 10e-9  # 10 ns, units are always S.I.
photon_data = dict(
    timestamps=timestamps,
    detectors=detectors,
    timestamps_specs={'timestamps_unit': timestamps_unit})
setup = dict(
    num_pixels = 2,                   # using 2 detectors
    num_spots = 1,                    # a single confoca excitation
    num_spectral_ch = 2,              # donor and acceptor detection 
    num_polarization_ch = 1,          # no polarization selection 
    num_split_ch = 1,                 # no beam splitter
    modulated_excitation = False,     # CW excitation, no modulation 
    excitation_alternated = [True],  # CW excitation, no modulation 
    lifetime = False,                 # no TCSPC in detection    
    excitation_wavelengths = [515e-9, 635e-9],         # List of excitation wavelenghts
    excitation_cw = [True],                    # List of booleans, True if wavelength is CW
    detection_wavelengths = [580e-9, 690e-9],  # Nominal center wavelength 
)
provenance = dict( 
    software='Custom Labview Software - Craggs Lab',
    filename=raw)
identity = dict(
    author= meta(3),
    author_affiliation="University of Sheffield") #CHANGE THIS WHERE NECESSARY
measurement_specs = dict(
    measurement_type = 'smFRET-usALEX',
    detectors_specs = {'spectral_ch1': [0],  # list of donor's detector IDs
                       'spectral_ch2': [1]},  # list of acceptor's detector IDs,
    alex_period = 10000,
    alex_offset = 0,
    alex_excitation_period1 = (D_ON, D_OFF),
    alex_excitation_period2 = (A_ON, A_OFF)
    )
photon_data['measurement_specs'] = measurement_specs
data = dict(
    description=meta(9),
    photon_data = photon_data,
    setup=setup,
    identity=identity,
    provenance=provenance
)
phc.hdf5.save_photon_hdf5(data, h5_fname=savename, overwrite=True)

# %% [code]
```
