## Supplementary material for "The smfBox: an open-source platform for single-molecule FRET": Jupyter Notebooks and data for accurate FRET: Correction Factor Finder Alpha-Delta.ipynb

```
# %% [markdown]
# Finding Alpha and Delta

This notebook is for extracting the alpha and delta accurate FRET correction factors from a dataset. You can load in data from a doubly labelled sample and select the donor only and acceptor only populations, or you can load in singly labelled samples and skip to that section.

Working from a doubly labelled sample has the disadvantage that only a minority of the bursts will be singly labelled. However, it is arguably more accurate to derive the correction factors from the same data set (or even just the same sample) as it guarantees that the fluorophores are in exactly the same environment in both the determination and application of the parameters.

# %% [markdown]
# Import packages

# %% [code]
from fretbursts import *
sns = init_notebook()
import lmfit
import phconvert
import os
from fretbursts.burstlib_ext import burst_search_and_gate

# %% [markdown]
# Name and Load in data

Name the data file and check it exists, note that it will look for files starting from the folder this notebook is in.

# %% [code]
filename = "definitiveset/1x.hdf5"
if os.path.isfile(filename):
    print("File found")
else:
    print("File not found, check file name is correct")

# %% [markdown]
Load in the file, we do not need to set correction factors yet.

You may get warnings that some parameters are not defined in the file, this is fine as they will be defined in this workbook anyway.

This cell also contains code to sort the timestamps as they are loaded. This is because the timestamps are not always saved in chronological order, and this can confuse some of FRETBursts functions.

# %% [code]
d = loader.photon_hdf5(filename)

for i in range(0, len(d.ph_times_t)): #sorting code
    indices = d.ph_times_t[i].argsort()
    d.ph_times_t[i], d.det_t[i] = d.ph_times_t[i][indices], d.det_t[i][indices]

# %% [markdown]
# Check alternation cycle is correct

We need to check that the ALEX parameters defined in the HDF5 file are appropriate for the laser cycle used in the experiment. If this is correct, the following histogram should look correct. It is a combined plot of every photon that arrives over the supplied alternation periods. 

# %% [code]
#d.add(det_donor_accept = (0, 1),
#    alex_period = 10000,
#    offset = 0,
#    D_ON = (0, 4500),
#    A_ON = (5000, 9500))
bpl.plot_alternation_hist(d)

# %% [markdown]
IF THE ABOVE HISTOGRAM LOOKS CORRECT: then run loader.alex_apply_period, which rewrites the time stamps into groups based on their excitation period. If you want to change the alternation period after this you will have to reload the data into FRETbursts.

IF THE ABOVE HISTOGRAM LOOKS WRONG: then the supplied alternation parameters do not match up to the alternation of the lasers in the data. This could be because the lasers were actually on a different alternation, or because the data set doesn't start at zero so is frame shifted etc.

In this case, you can un-hash the code below and alter the parameters manually.

# %% [code]
#d.add(det_donor_accept = (0, 1),
#    alex_period = 10000,
#    offset = 0,
#    D_ON = (0, 4500),
#    A_ON = (5000, 9500))

# %% [code]
loader.alex_apply_period(d)
time = d.time_max
print('Total data time = %s'%time)

# %% [markdown]
The following will plot a time trace of the first second of your experiment. You can change "start, stop =" to look through the trace.

# %% [code]
start, stop = 0, 1
dplot(d, timetrace, binwidth=1e-3, tmin=0, tmax=15, figsize=(8,5))
plt.xlim(start, stop);
plt.ylim(-45,45);

plt.ylabel(" Photons/ms in APD0       Photons/ms in APD1");

# %% [markdown]
# Background Estimation

# %% [markdown]
Background estimation works by plotting log of photons by the delay between them, assuming a poisson distribution of photon arrivals and fitting a line. The plot will contain single molecule bursts however, so a threshold (in microseconds) has to be defined where the fit begins. Choosing the right threshold is a trade off between correctly ignoring the burst photons (accuracy) and getting enough photons to calculate the background (precision). 

"time_s=" will change how regularly the background is recalculated

# %% [code]
threshold = 1500
d.calc_bg(bg.exp_fit, time_s=300, tail_min_us=(threshold))
dplot(d, hist_bg, show_fit=True)

# %% [markdown]
# Burst Searching

M = the number of photons to look at at any time
F = the number of times higher than the background the photon rate must be to be considered a burst
L = the number of photons that must be in a burst to be considered a burst

The burst selection then selects only bursts that are size th1 or higher, and lower than th2. add_naa=True ensures acceptor photons under acceptor excitation are encluded in the search.

Here we use an APBS, then select from an all photon count so that we do not bias in S.

# %% [code]
d.burst_search(L=10, m=10, F=45)
ds = d.select_bursts(select_bursts.size, add_naa=True, th1=50)

# %% [markdown]
now we can start plotting and selecting populations

# %% [code]
plt.rcParams['axes.facecolor'] = 'white'
grids = False
colourscheme = 'viridis_r'
g=alex_jointplot(ds, cmap=colourscheme, marginal_color=20, vmax_fret=True) 

# %% [markdown]
# Alpha

Alpha, or the leakage factor, describes the amount of photons emitted by the donor which are detected by the acceptor APD (relative to the donor APD), as a consequence of the red tail of the donor emission spectrum. It can be determined from the apparent FRET efficiency of the donor only population

# %% [markdown]
Now we select the donor only population by drawing a box.

# %% [code]
roi= dict(E1=-0.2, E2=0.5, S1=0.95, S2=1.1, rect=True)
donoronly = ds.select_bursts(select_bursts.ES, **roi)
g = alex_jointplot(donoronly, cmap=colourscheme, marginal_color=20, vmax_fret=False)
bpl.plot_ES_selection(g.ax_joint, **roi);

# %% [markdown]
This will fit a single gaussian to the filtered E, and convert the mean position to an alpha value, the leakage factor

# %% [code]
model = mfit.factory_gaussian()
donoronly.E_fitter.fit_histogram(model=model, verbose=False, pdf=False)
dplot(donoronly, hist_fret, binwidth=0.025, show_model=True, pdf=False, show_fit_value=True, fit_from="center");
params = donoronly.E_fitter.params
Efit = params.to_dict()
Centre = Efit['center']
Edonly=Centre[0]
Edonly
alpha=Edonly/(1-Edonly)
print("alpha =", alpha)

# %% [markdown]
# Delta

Delta, or the direct excitation factor, describes the relative emission of the donor and acceptor under excitation of the donor laser, as a consequence of the blue tail of the acceptor absorption spectrum. It can be determined from the apparent stoichiometry of the acceptor only population.

# %% [code]
roi= dict(E1=-0.2, E2=1.2, S1=-0.2, S2=0.175, rect=True)
acceptoronly = ds.select_bursts(select_bursts.ES, **roi)
g = alex_jointplot(acceptoronly, cmap=colourscheme, marginal_color=20, vmax_fret=False)
bpl.plot_ES_selection(g.ax_joint, **roi);

# %% [markdown]
This will extract delta, the direct excitation factor from the S histogram

# %% [code]
model = mfit.factory_gaussian()
acceptoronly.S_fitter.fit_histogram(model=model, verbose=False, pdf=False)
dplot(acceptoronly, hist_S, binwidth=0.025, show_model=True, pdf=False, show_fit_value=True, fit_from="center");
acceptoronly.S_fitter.params
params = acceptoronly.S_fitter.params
Sfit = params.to_dict()
Centre = Sfit['center']
Saonly=Centre[0]
delta=Saonly/(1-Saonly)
print("delta =", delta)

# %% [code]
print("Alpha (lk factor)      = ", round(alpha, 5))
print("Delta (dir. ex factor) = ", round(delta, 5))

# %% [code]
```
