## Supplementary material for "The smfBox: an open-source platform for single-molecule FRET": Jupyter Notebooks and data for accurate FRET: Correction Factor Finder Gamma-Beta.ipynb

```
# %% [markdown]
# Finding Gamma and Beta

This notebook is for calculating the gamma and beta correction factors from multiple loaded datasets. It is possible to find these parameters from a single data set so long as the distribution of apparent FRET efficiencies is dominated by a breadth of real underlying distances, however in this case we load multiple data sets each containing a single FRET state.

Gamma describes the inequality between how well the donor and acceptor fluorophores can be detected by your system, whereas beta describes the inequality between how well they can be excited. They are both found by an equation which describes the slope and height (in S) of the doubly labelled population. 

You should find the alpha and delta factors first before this step.

# %% [markdown]
# Import packages

# %% [code]
from fretbursts import *
sns = init_notebook()
import lmfit
import phconvert
import os
from fretbursts.burstlib_ext import burst_search_and_gate

# %% [markdown]
# Name and Load in data

Name the data files, note that it will look for files starting from the folder this notebook is in.

# %% [code]
files = ["definitiveset/1a1.hdf5", "definitiveset/1a2.hdf5",
         "definitiveset/1b1.hdf5", "definitiveset/1b2.hdf5",
         "definitiveset/1c1.hdf5", "definitiveset/1c2.hdf5"]

# %% [markdown]
Set the spectral cross talk parameters, load them, and sort them

# %% [code]
alpha = 0.081
delta = 0.076

datasets = []
for file in files:
    datasets.append(loader.photon_hdf5(file))
for dataset in datasets:
    dataset.leakage = alpha
    dataset.dir_ex = delta
    for i in range(0, len(dataset.ph_times_t)): #sorting code
        indices = dataset.ph_times_t[i].argsort()
        dataset.ph_times_t[i], dataset.det_t[i] = dataset.ph_times_t[i][indices], dataset.det_t[i][indices]

# %% [code]
datasets

# %% [markdown]
# Apply alternation cycle

# %% [code]
for dataset in datasets:
    #dataset.add(det_donor_accept = (0, 1),
    #alex_period = 10000,
    #offset = 0,
    #D_ON = (0, 4500),
    #A_ON = (5000, 9500))
    loader.alex_apply_period(dataset)

# %% [markdown]
# Background Estimation

# %% [markdown]
We background correct all datasets as before

# %% [code]
threshold = 1500
recalctime = 300
for dataset in datasets:
    dataset.calc_bg(bg.exp_fit, time_s = recalctime, tail_min_us=(threshold))
    dplot(dataset, hist_bg, show_fit=True)

# %% [markdown]
Now let's find bursts and plot them. We will use a dual channel burst search this time as we are looking for strictly doubly labelled bursts.

# %% [code]
colourscheme= "viridis_r"
burstsets = []
for dataset in datasets:
    dataset = burst_search_and_gate(dataset, F=45, m=10, mute=True)
    dataset = dataset.select_bursts(select_bursts.size, th1=50)
    burstsets.append(dataset)
for burstset in burstsets:
    alex_jointplot(burstset, cmap=colourscheme, marginal_color=20, vmax_fret=False)

# %% [markdown]
Now let's extract the E and S positions from these plots with gaussian fits. We could plot the fitted curves, but here the graphs are not shown to save space.

# %% [code]
def ESgauss(dataset):
    dataset.E_fitter.fit_histogram(model=mfit.factory_gaussian(), verbose=False, pdf=False)
    params = dataset.E_fitter.params
    Efit = params.to_dict()
    E = Efit['center'][0]
    dataset.S_fitter.fit_histogram(model=mfit.factory_gaussian(), verbose=False, pdf=False)
    params = dataset.S_fitter.params
    Sfit = params.to_dict()
    S = Sfit['center'][0]
    return(E, S)
x = []
y = []
for dataset in burstsets:
    E, S = ESgauss(dataset)
    x.append(E)
    y.append(S)    

# %% [markdown]
Now we can fit to the equation:

    S = 1/(1+bg+(1-g)*b*E)

# %% [code]
model = lmfit.models.ExpressionModel('1 / ( 1 + b * g + ( 1 - g ) * b * E )',
                             independent_vars=["E"], nan_policy = "omit")
params = model.make_params(b=0.5, g=1.0)
fit = model.fit(y, params, E=E)
fit.plot_fit(0)
print(fit.fit_report())

# %% [code]
pars = fit.params
beta = pars['b']
beta=beta.value
gamma = pars['g']
gamma=gamma.value
print("gamma = ", gamma)
print("beta = ", beta)

# %% [code]
```
