## Supplementary figures and images for "The smfBox: an open-source platform for single-molecule FRET"

### Technical_Drawing_smfScope_APD_Cover.png

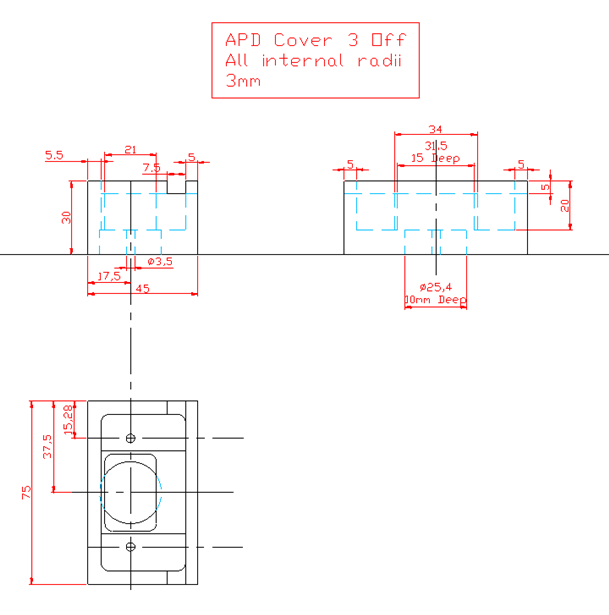

### Technical_Drawing_smfScope_APD_Mount.png

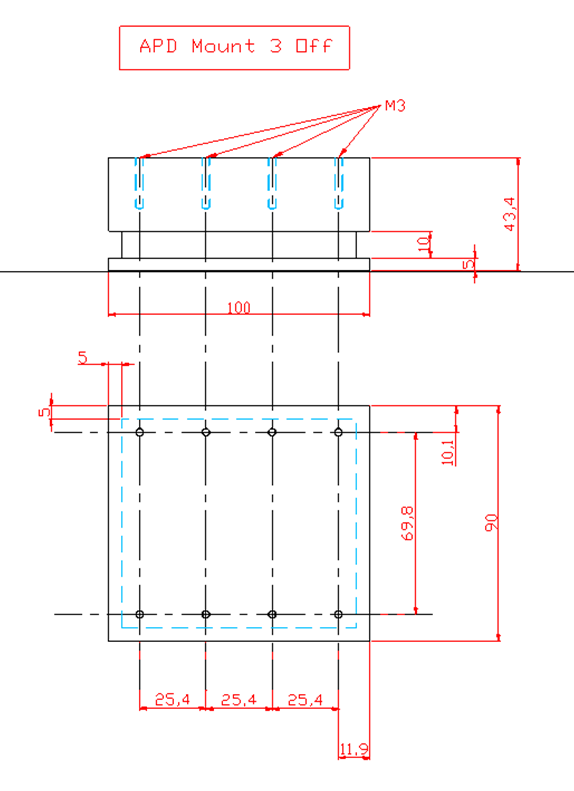

### Technical_Drawing_smfScope_Box_Back_Plate.png

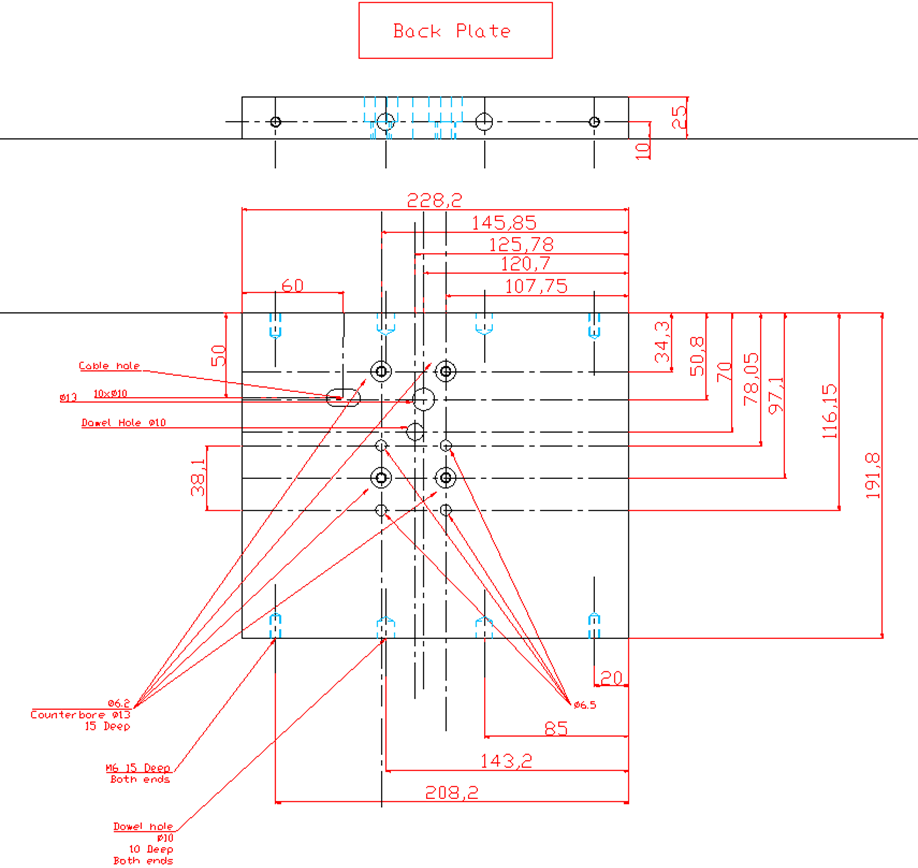

### Technical_Drawing_smfScope_Box_Base_Plate.png

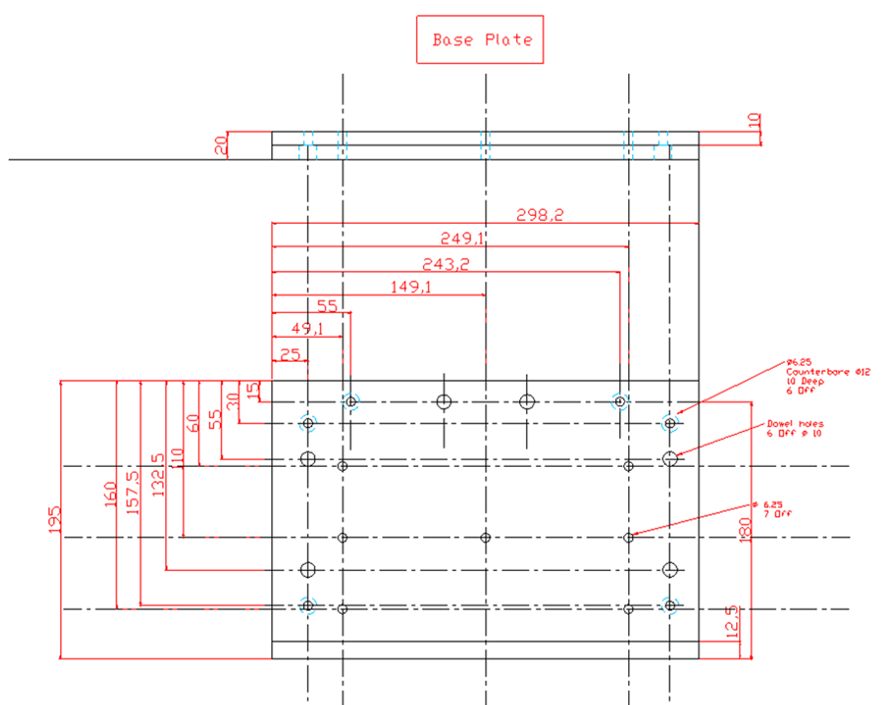

### Technical_Drawing_smfScope_Box_Block.png

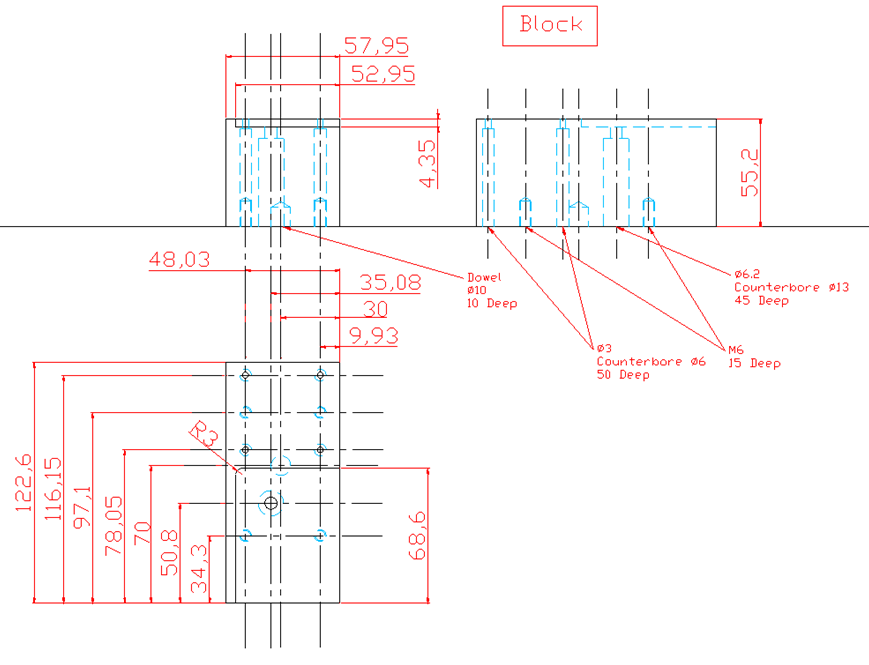

### Technical_Drawing_smfScope_Box_Front_Plate.png

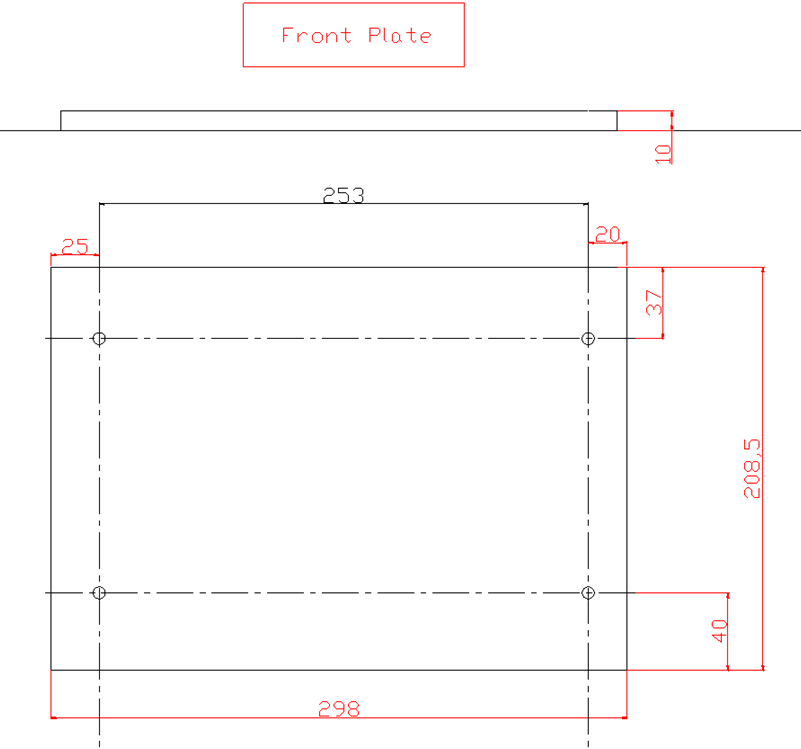

### Technical_Drawing_smfScope_Box_Laser-In_Plate.png

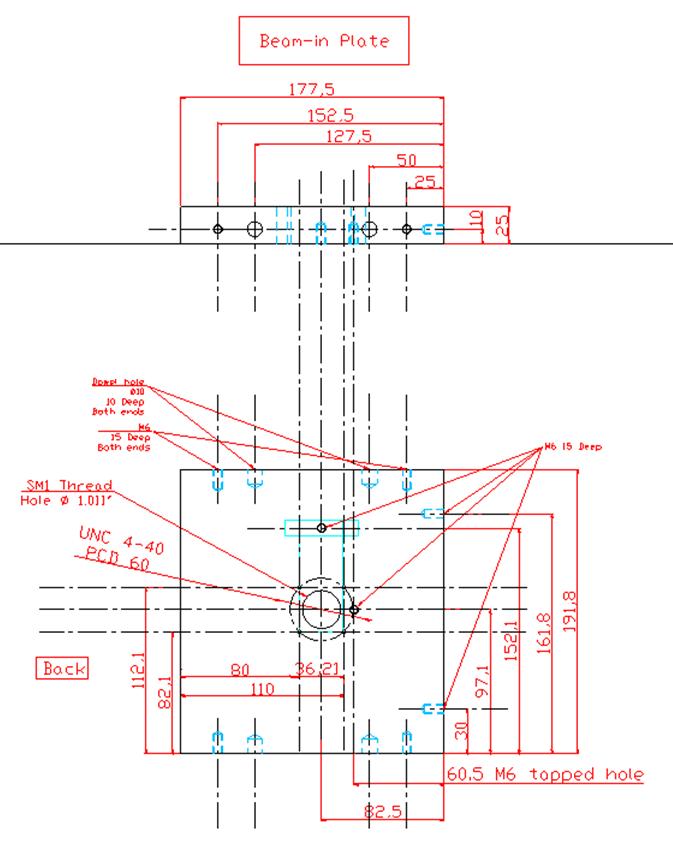

### Technical_Drawing_smfScope_Box_Laser-Out_Plate.png

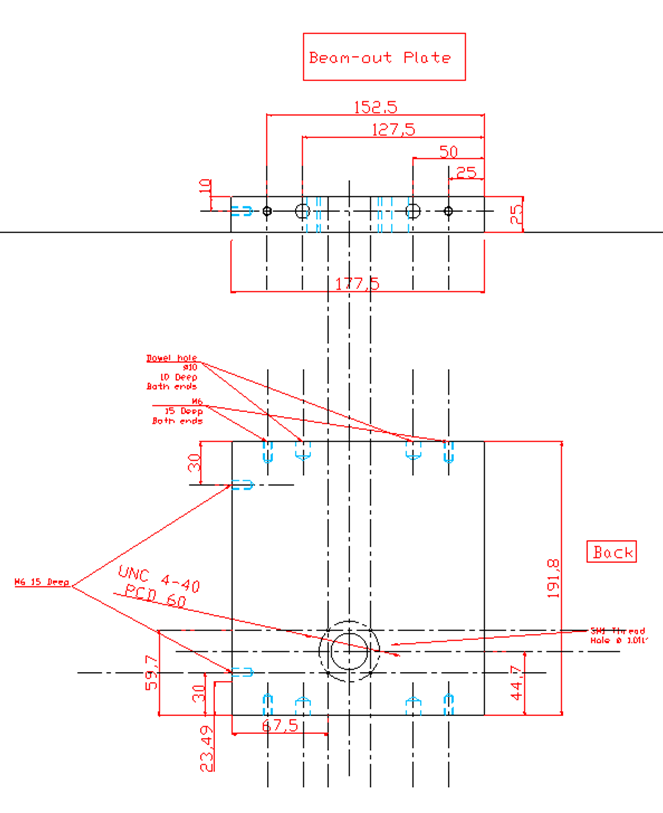

### Technical_Drawing_smfScope_Box_Lid.png

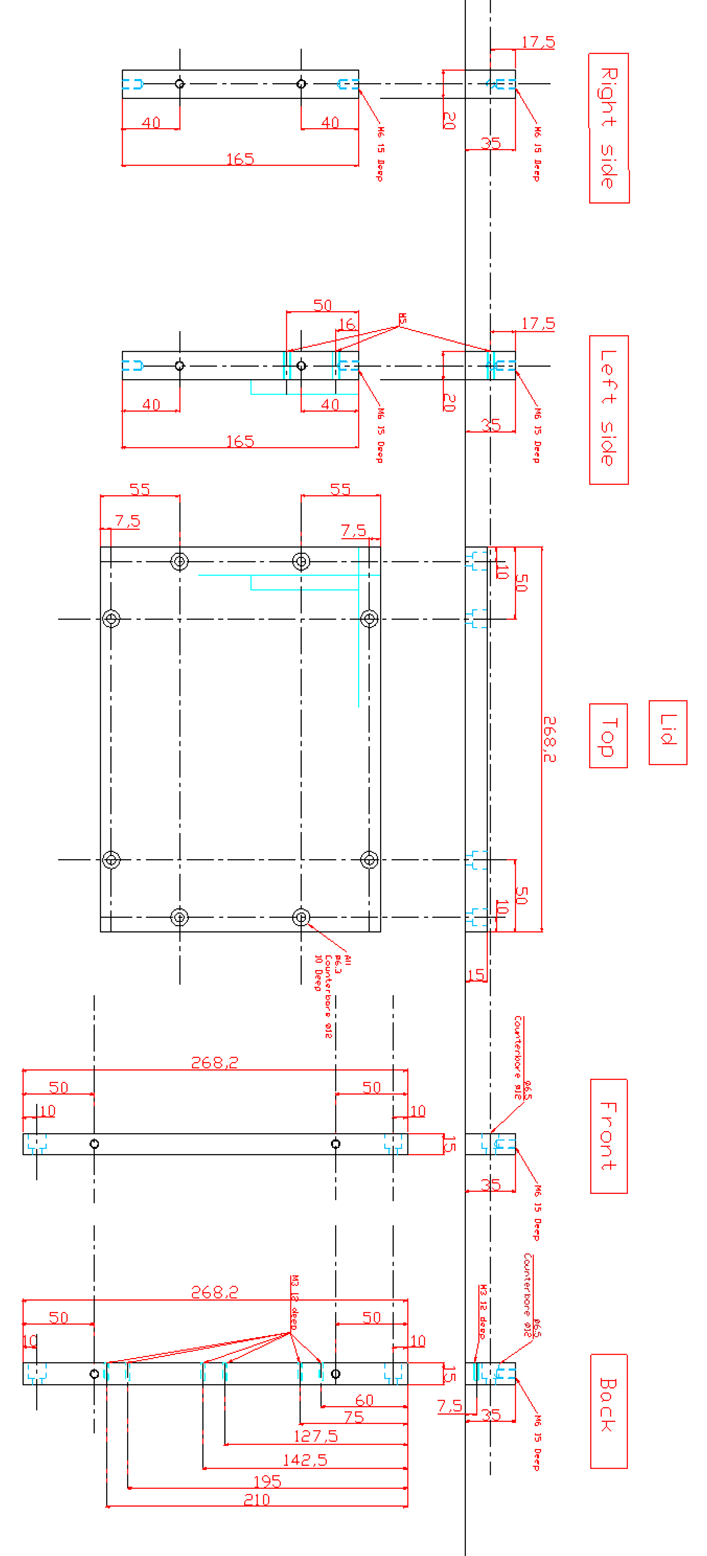

### Technical_Drawing_smfScope_Box_Shutter.png

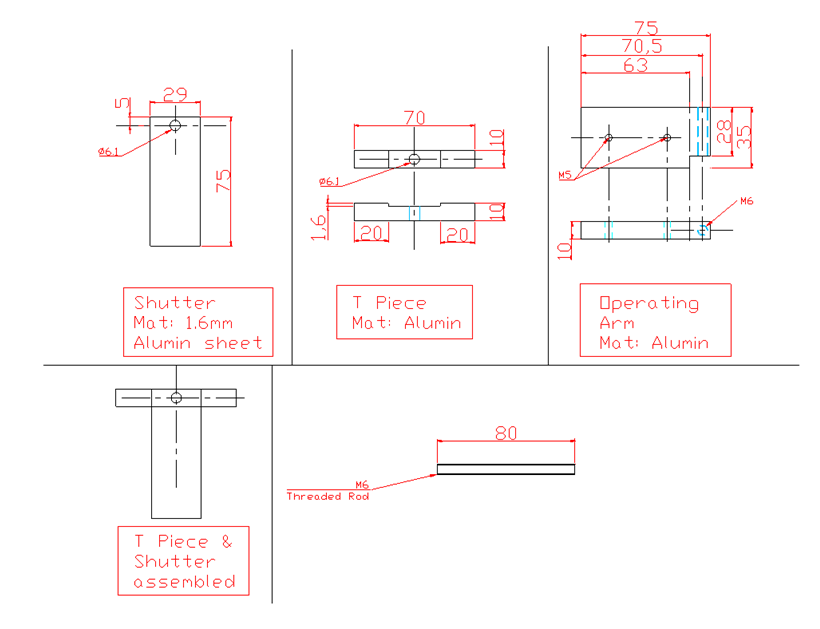

### Technical_Drawing_smfScope_Box_Shutter_Assembly.png

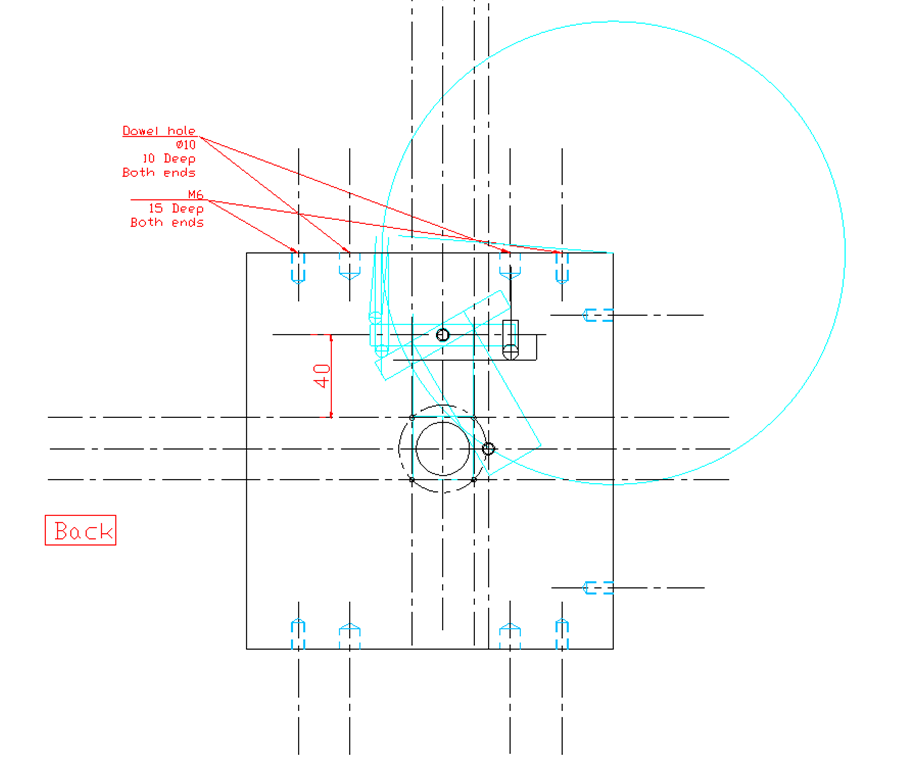

### Technical_Drawing_smfScope_Box_Top_Plate.png

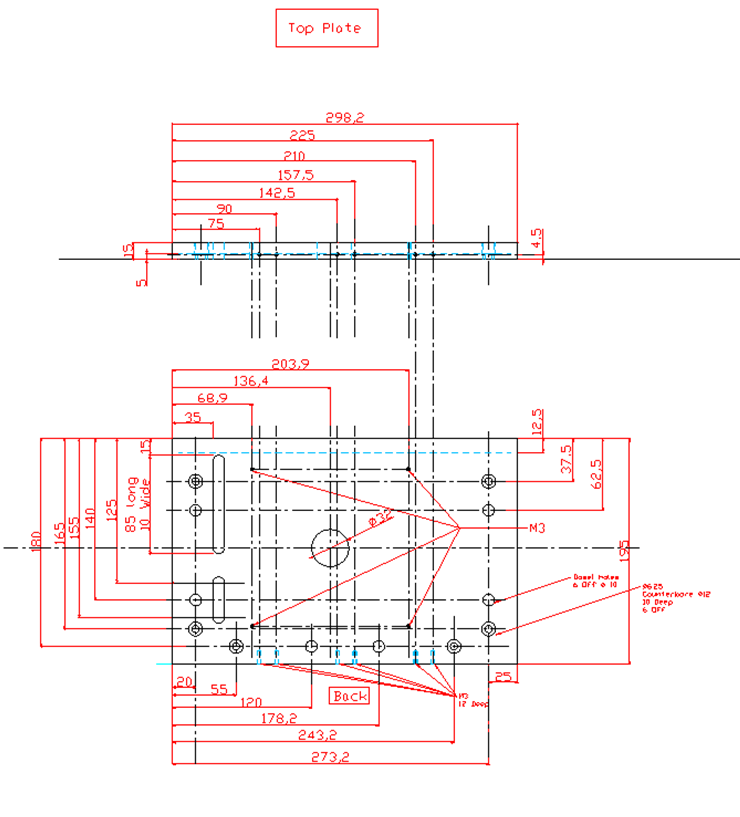

### Technical_Drawing_smfScope_Shutter_Box_Assembly_Top_Plate.png

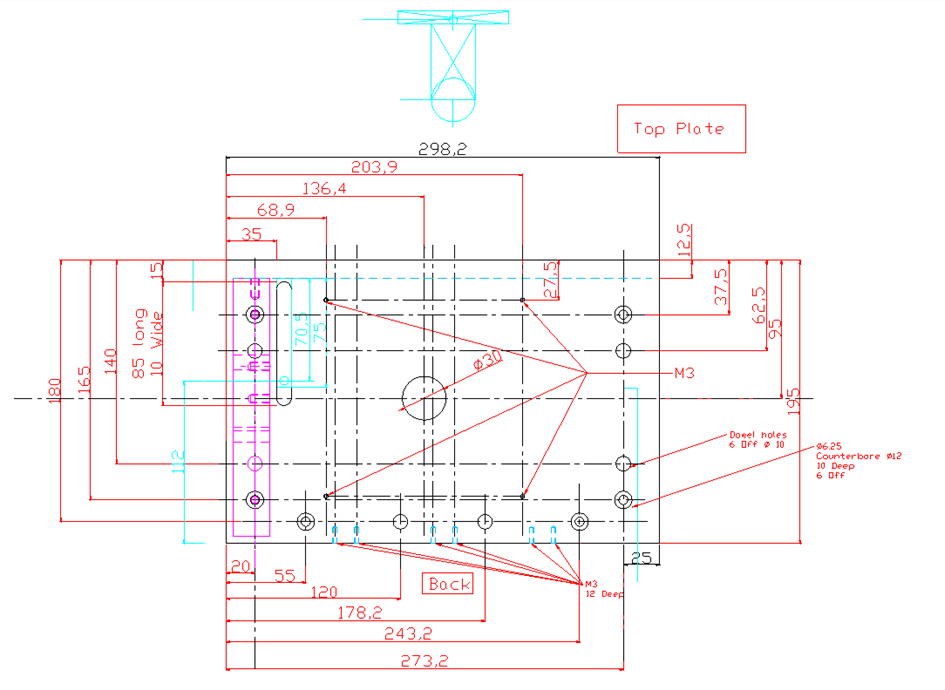

### Technical_Drawing_smfScope_Stage.png

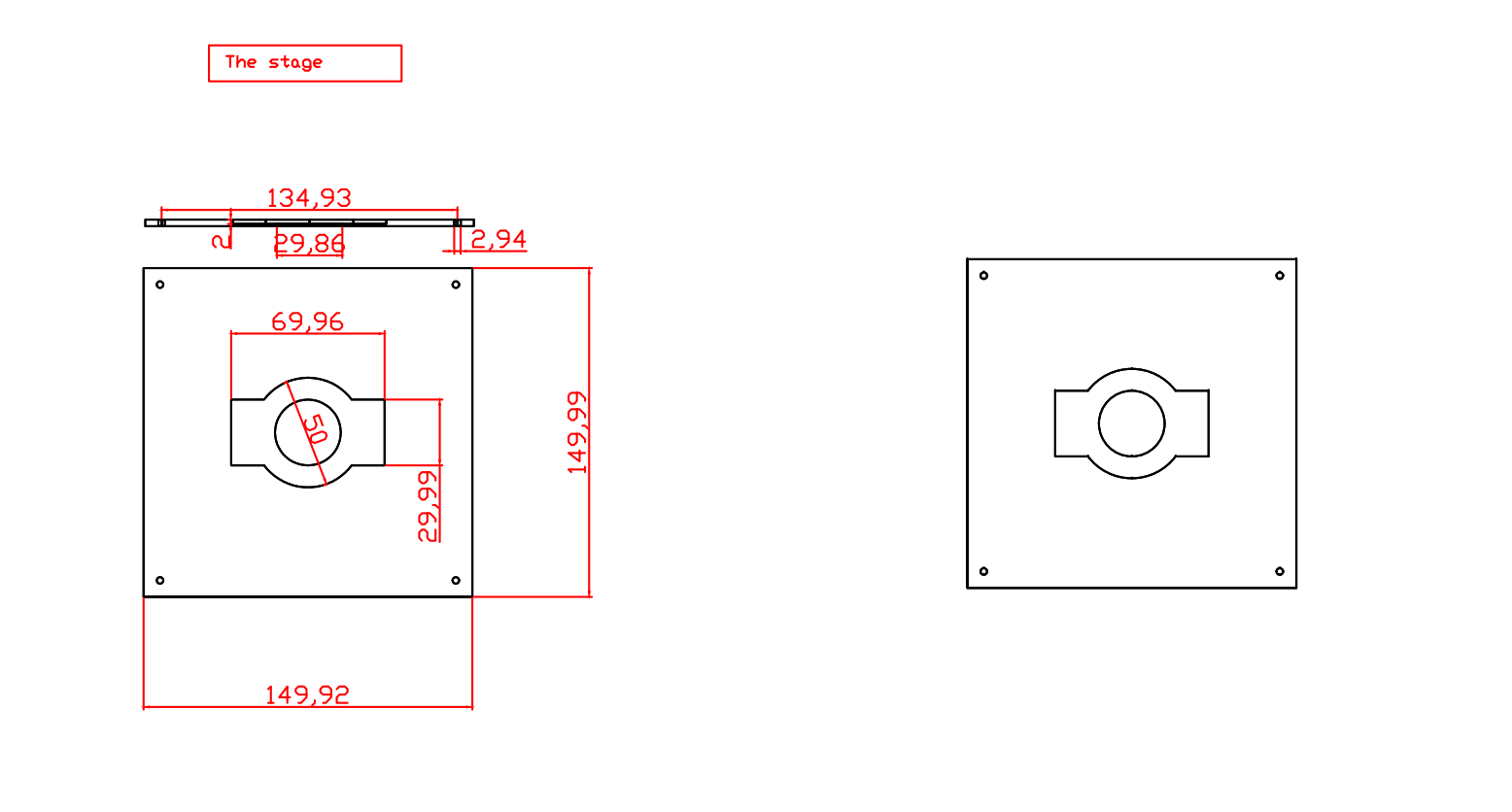
