## Supplementary material for "The smfBox: an open-source platform for single-molecule FRET": Labview operational software: Microscope Alignment.ini

```
[Microscope Alignment]
server.app.propertiesEnabled=True
server.tcp.paranoid=True
server.tcp.serviceName="My Computer/VI Server"
server.vi.callsEnabled=True
server.vi.propertiesEnabled=True
WebServer.TcpAccess="c+*"
WebServer.ViAccess="+*"
DebugServerEnabled=False
DebugServerWaitOnLaunch=False
```
